# A *P2rx7* passenger mutation affects the vitality and function of immune cells in P2X4ko and other transgenic mice

**DOI:** 10.1101/2020.06.10.139410

**Authors:** Marco Er-Lukowiak, Yinghui Duan, Francois Rassendren, Lauriane Ulmann, Annette Nicke, Friederike Ufer, Manuel A. Friese, Friedrich Koch-Nolte, Tim Magnus, Björn Rissiek

**Affiliations:** Department of Neurology, University Medical Centre Hamburg-Eppendorf, Hamburg, Germany; IGF, Univ. Montpellier, CNRS, INSERM, Montpellier, France; LabEx ICST, Montpellier, France; Institut de Génomique Fonctionnelle, CNRS UMR5203, Montpellier, France; Walther Straub Institute for Pharmacology and Toxicology, Ludwig-Maximilians-Universität München, Munich, Germany; Institute of Neuroimmunology und Multiple Sclerosis (INIMS), University Medical Centre Hamburg-Eppendorf, Hamburg, Germany; Institute of Immunology, University Medical Centre Hamburg-Eppendorf, Hamburg, Germany

**Author notes:** Equally contributing senior authors (Friedrich Koch-Nolte, Tim Magnus, Björn Rissiek). To whom correspondence should be addressed: Dr. Björn Rissiek, University Medical Center Hamburg-Eppendorf, Department of Neurology, Martinistraße 52, 20246 Hamburg-Eppendorf, Germany.

## Abstract

Among laboratory mouse strains many genes are differentially expressed in the same cell population. As consequence, gene targeting in 129-derived embryonic stem cells (ESCs) and backcrossing the modified mice onto the C57BL/6 (B6) background can introduce passenger mutations in the close proximity of the targeted gene. Here, we demonstrate that several 129-originating transgenic mice in which *P2rx7*-neighboring genes were targeted carry a *P2rx7* passenger mutation that affects the vitality and function of T cells. By the example of *P2rx4^tm1Rass^* we demonstrate that CD4^+^ and CD8^+^ T cells derived from these mice express higher levels of P2X7 when compared to corresponding cell populations in B6-WT mice. The increased T cell sensitivity towards the P2X7 activators adenosine triphosphate (ATP) and nicotinamide adenine dinucleotide (NAD^+^) rendered these cells more vulnerable towards NAD-induced cell death (NICD) compared to their B6-WT counterparts. The enhanced NICD sensitivity significantly affected the outcome of functional assays e.g. cytokine production and cell migration. For P2rx4^tm1Rass^, we demonstrate that the expression of P2X7 is diminished in several innate immune cell populations, possibly as a side effect of *P2rx4* targeting, and independent of the *P2rx7* passenger mutation. These results need to be considered when working with P2rx4^tm1Rass^ mice or other 129-based transgenic strains that target *P2rx7* neighboring genes and might have implications for other mouse models.

## Introduction

In the pre-CRISPR-Cas9 era, gene targeting, in order to generate “knockout” or “knockin” mice, was often conducted in embryonic stem cells (ESCs) derived from 129-originating mouse strains. The obtained transgenic mice were then backcrossed on C57BL/6, the most widely used strain in immunological research. Due to genetic linkage, however, flanking regions of the targeted gene are unlikely to be exchanged by backcrossing and the likelihood of exchange decreases the closer these regions are located to the targeted gene (Lusis et al., 2007). Genetic distances are often given in centimorgans (cM), with 1 cM distance representing the equivalent to 1% probability of two loci being separated during homologous recombination in the context of meiosis. In the mouse, 1 cM is, on average, equivalent to 2,000 kilobases, however, the rate of equivalence can vary greatly due to numerous factors (Silver, 1995). This implicates a probability of about 90% that upon backcrossing for 10 generations, a flanking region with a distance of 1 cM to the targeted gene is still of donor origin (Berghe et al., 2015; Lusis et al., 2007). Gene expression can vary in cells from different laboratory mouse strains and some mouse strains even constitute “natural knockouts” for certain genes e.g. if mutations or SNPs introduce premature STOP codons (Mostafavi et al., 2014). If such a differentially expressed or inactive gene is in close proximity of a targeted gene in 129 ESCs, backcrossing to C57BL/6 will introduce an experimental bias, since the differentially expressed/inactive gene most likely will be retained. A prominent example is the 129-derived Casp1 knockout mouse (e.g. Casp1^tm1Flvw^). The 129 mouse strains carry a defective *Casp11* gene in the close proximity of *Casp1* and the strong resistance to lethal lipopolysaccharide (LPS) injection found in Casp1^-/-^ mice was mainly due to the defective *Casp11* neighboring gene that was carried along as passenger mutation (Kayagaki et al., 2011). A later study could show that targeting of other *Casp11* neighboring genes, such as *Panx1*, also led to a resistance towards lethal LPS injection, if ESCs were derived from 129 mice, but not when B6-derived ESCs were used (Berghe et al., 2015). These two examples illustrate the potential impact of 129-derived passenger mutations and possible consequences, if they remain undetected.

P2X7 is a homotrimeric, adenosine triphosphate (ATP)-gated ion channel expressed on the cell surface of many immune cells (Bartlett et al., 2014). During tissue damage and inflammation ATP is released into the extracellular space where it serves as damage associated molecule (DAMP). ATP-induced activation of P2X7 in macrophages and microglia triggers the formation of the NACHT, LRR and PYD domains-containing protein 3 (NLRP3) inflammasome, consisting of NLRP3, the adaptor protein Apoptosis-Associated Speck-Like Protein Containing CARD (ASC) and caspase 1. The NLRP3 inflammasome catalyzes the processing of pro-interleukin 1 beta (pro-IL-1β) into its active form IL-1β (Ferrari et al., 1997; 2006). In the mouse, certain T cell populations have been identified as “high P2X7” expressers: these are regulatory T cells (Treg), cytotoxic lymphocytes (CTL) from the lamina propria, natural killer T cells (NKT), follicular helper T cells (Tfh) and tissue-resident memory T cells (Trm) (Aswad et al., 2005; da Silva et al., 2018; Heiss et al., 2008; Hubert et al., 2010; Kawamura et al., 2006; Proietti et al., 2014; Rissiek et al., 2014; 2018b). Autocrine ATP release followed by activation of P2X7 and other P2X receptors plays a role in various T cell functions, including in IL-2 secretion, metabolic fitness and cell migration (da Silva et al., 2018; Ledderose et al., 2018; Rissiek et al., 2015). High concentrations of ATP and prolonged activation of P2X7 on T cells, however, is a strong trigger of T cell death (Scheuplein et al., 2009). In contrast to human T cells, mouse T cells exhibit an alternative way of P2X7 activation, which is triggered by ecto-ADP-ribosyltransferase C2.2 (ARTC2.2). ARTC2.2 utilizes nicotinamide adenine dinucleotide (NAD^+^), which is also released as DAMP during tissue damage, to ADP-ribosylate a variety of cell surface proteins such as CD25 (Teege et al., 2015), CD8β (Lischke et al., 2013), FcγR1, FcγR2b (Rissiek et al., 2017) and P2X7 (Seman et al., 2003). ADP-ribosylation of arginine 125 in the extracellular loop of P2X7 (Adriouch et al., 2008) acts like a covalently bound P2X7 agonist (Schwarz et al., 2009). Of note, 10-fold lower concentrations of NAD^+^ compared to ATP are needed to activate T cell P2X7 receptors and its ADP-ribosylation ultimately leads to NICD (Seman et al., 2003). Our own studies revealed that NAD^+^ is also released during preparation of primary T cells from mouse organs such as spleen and lymph nodes (Rissiek et al., 2014; Scheuplein et al., 2009). Interestingly, ARTC2.2 is active at 4°C, however ADP-ribosylation-mediated gating of P2X7 needs temperatures above 30°C (Scheuplein et al., 2009). This implicates that freshly on ice prepared T cells appear vital, but incubation at 37°C, can trigger cell death on T cells that co-express high level of ARTC2.2 and P2X7 skewing the results of cytokine secretion assays as has been demonstrated for Trm, NKT and Tfh (Borges da Silva et al., 2019; Georgiev et al., 2018; Rissiek et al., 2018b). The problem of P2X7 ADP-ribosylation during cell preparation can be solved by injecting an ARTC2.2 blocking nanobody (clone s+16a) into the mice 30 min before harvesting the mouse organs (Hubert et al., 2010; Koch-Nolte et al., 2007; Rissiek et al., 2014).

In the mouse, the *P2rx7* gene is located on chromosome 5 with 95 other characterized, protein-coding genes being within a distance of 2 megabases (Mb) upstream and downstream of *P2rx7* **(Table 1)**. For *P2rx7*, a single nucleotide polymorphism (rs48804829) introduces a change of proline to leucin at amino acid position 451 in the cytosolic C-terminal tail of P2X7, which is associated with reduced ATP-mediated pore formation (Adriouch et al., 2002). The 129 and Balb/c mouse strains harbor the P2X7^451P^ variant, whereas C57BL/6 (B6) mice express the loss-of-function P2X7^451L^ variant. Studies revealed that P2X7^451P^ and P2X7^451L^ differ in their capacity to induce the formation of a P2X7-associated membrane pore (Adriouch et al., 2002) but not in P2X7-triggered calcium influx (Le Stunff et al., 2004; Sorge et al., 2012). No comparative P2X7 expression studies among mouse strains that express P2X7^451P^ and P2X7^451L^ have been performed. In this study, we analyzed P2X7 expression on T cell populations from different mouse strains. We show that CTL and helper T cells (Th) from strains that harbor P2X7^451P^ express much higher P2X7 levels than cells derived from P2X7^451L^ strains. As 129 mice express P2X7^451P^ whereas B6 mice express P2X7^451L^, we hypothesized that a 129-derived *P2rx7* gene could introduce an experimental bias when passed along as passenger mutation in congenic mice. Here we demonstrate that in P2rx4^tm1Rass^ mice Th and CTLs are much more prone to NICD compared to their wildtype (WT) counterparts. As consequence, these T cells appear less potent in terms of cytokine production and migration. However, when ARTC2.2 is blocked during cell preparation, the functional deficits of B6-P2X4ko T cells vanish. We identify other mouse strains with mutations in neighboring genes that also carry the 129-derived *P2rx7* passenger mutation. Our study emphasizes the importance of checking the ESC origin of transgenic mice and to analyze them for passenger mutations in order to prevent misinterpretation of experimental results.

## Material and Methods

### Mice

Mice on the C57BL/6 and Balb/c background were used in this study. P2rx4^tm1Rasss^ on the B6 background were obtained from the lab of Francois Rassendren. Balb/c-P2X4^-^ko mice were generated by backcrossing B6-P2X4ko mice with Balb/c-WT mice for 13 generations. In some experiments cells from B6-P2X7ko mice (P2rx7^tm1Gab^) were used as negative control for P2X7 cell surface staining. Hvcn1^Gt(RRRN293)Byg^ on the B6 background were obtained from the lab of David Clapham. All mouse experiments were approved by the responsible regulatory committee (Hamburger Beho□rde fu□r Gesundheit und Verbraucherschutz, Veterina□rwesen/Lebensmittelsicherheit, ORG722, ORG983, G12/130). All experiments were performed according to the relevant guidelines and regulations.

### Antibodies and flow cytometry

For FACS analysis, the following antibodies were used: anti-CD3-Bv421-(clone 17A2, Biolegend), anti-CD4-Bv421 (clone RM 4-5, eBioscience), anti-CD8-FITC (clone 53-6.7, Biolegend), anti-CD25 PE (clone PC61, Biolegend), anti-CD27-APC (clone LG.3A10, Biolegend), anti-CD11b-FITC (clone M1/70, Biolegend), anti-CD11b-Bv421 (clone M1/70, Biolegend), anti-FceRIα-PE (MAR-1, Biolegend), anti-F4/80-PE (BM8, Biolegend), anti-CD45-APC/Cy7 (30-F11, Biolegend), anti-CD107a-FITC (1D4B, Biolegend) and anti-P2X7-AF647 (clone Hano44, UKE) (Adriouch et al., 2005). Flow cytometric analyses were performed on a BD Fortessa (Beckton Dickinson) or a BD FACS CantoII (Beckton Dickinson).

### Preparation of immune cells

The isolation of immune cells was performed strictly at 4°C on ice. Spleen and peripheral lymph nodes were mashed through a cell strainer (50 mL falcon strainer, 70 μm, GBO) using a syringe piston. Single cell suspension was kept in FACS buffer containing 1 mM EDTA (Sigma) and 0.1 % bovine serum albumin (Sigma). Erythrocytes were lyzed using an ACK lysis buffer (155 mM NH_4_Cl, 10 mM KHCO_3_, 0.1 mM EDTA, pH 7.2). For microglia preparation, brain tissue was digested with collagenase (1 mg/ml) and DNase (0.1 mg/ml) at 37°C for 30 min. The generated cell suspension was filtered through a 70 μm cell strainer and centrifuged for 5 min at 300 g. Microglia were separated from debris by resuspending the pellet in 5 ml 33% percoll solution (GE Healthcare). Mast cells and macrophages were obtained from the peritoneal cavity by lavage with 5 ml cold PBS + EDTA (2 mM, Gibco). Cells were resuspended in RPMI medium containing 10% FCS.

For some T cell experiments, mice were injected (i.v.) with 50 μg of the ARTC2.2-blocking nanobody s+16a 30 minutes prior to sacrificing the mice in order to prevent ADP-ribosylation of P2X7 on T cells during cell preparation.

### Generation of bone marrow-derived innate immune cells

Bone marrow was obtained from mouse femurs. The distal and proximal ends of the femurs were cut off and the bone marrow was flushed out with cold PBS using a syringe. Total cells were count (Neubauer chamber), washed once with sterile PBS. A total amount of 6 x 10^6^ or 3 x 10^6^ bone marrow cells were cultured in 10 ml RPMI complete medium (glutamine 20 μM, 10 % FCS, β-mercaptoethanol 50 μM, gentamycin 50 μg/ml) supplemented with 10 ng/ml M-CSF (Biolegend) for BMM generation or 10 ng/ml GM-CSF (Biolegend) for BMDC generation. At day 3, 5 ml BMM and 10 ml BMDC medium was added to the corresponding cells. At day 6, total BMM medium was replaced with 10 ml fresh one while half of the BMDC medium was replaced with fresh one. After 7 days, the cells were ready for experimental use.

For the generation of BMMCs, 10 x 10^6^ bone marrow cells were cultured in RPMI 1640 medium containing + L-glutamine, 10 mM HEPES, 10% FCS, 50 μM gentamicin, 50 μM β-mercaptoethanol, 70 μM non-essential amino acids, 1 mM sodium pyruvate (Sigma) and recombinant IL-3 (10 ng/ml, Biolegend). BMMCs were transferred into a new petri dish with new BMMC medium every 3-4 days. BMMCs were ready to use after one month.

### Quantitative real-time PCR

RNA was extracted from FACS sorted immune cells using the RNeasy Plus Mini Kit (Qiagen) followed by cDNA synthesis using the Maxima First Strand cDNA Synthesis Kit (Thermo Fisher Scientific) as recommended by the respective supplier. RT-qPCR was performed on a Lightcycler 96 (Roche). The following Taqman probes (Thermo Scientific) were used for experiments: *Sdha* (Mm01352366_m1), *P2rx7* (Mm00440582_m1), *Camkk2 (Mm00520236), Atp2a2 (Mm01201431), Kdm2b (Mm01194587), Anapc5 (Mm00508066), and Anapc7 (Mm00517286)*.

### P2rx7 SNP sequencing

Sequencing of a region flanking SNP rs48804829 in the *P2rx7* gene was performed using the primers P2×7_P451L_forw (gggaaaagtctgcaagttgtc) and P2×7_P451L_rev (gaagagcttggaggtggtg). The PCR product was purified with the PCR clean-up gel extraction kit (Macherey-Nagel) and send to Eurofins, Germany, for sequencing.

### Monitoring P2X7 induced cell death on T cells

T cells were isolated by flow cytometric cell sorting. 5 x 10^4^ cell were resuspended in 400 μl complete RPMI medium containing propidium iodide PI (2.5 μg/ml, ImmunoChemistry Technologies). Half of the sample was left at 4 °C and the other half was incubated for 2 hours at 37 °C. Cell vitality was analyzed directly after incubation by flow cytometry.

### Monitoring P2X7 shedding of CD27 on T cells

For endpoint analyses, purified splenocytes from WT and P2X4ko mice were stained with lineage markers and anti-CD27 for 30 min, then washed and WT or P2X4ko cells were labeled with eFluor^670^. The labeled and unlabeled samples were mixed in a 1:1 ratio and aliquots were subjected to ATP dose response analyses(16-500 μM ATP). For this, cells were incubated in the presence of ATP at 37 °C for 15 minutes, samples without ATP were incubated at 4°C and 37°C as controls. Loss of CD27 from the cell surface was analyzed by flow cytometry.

For the real-time CD27 measurements, WT or P2X4ko cells were labeled with eFluor^670^. The labeled and unlabeled samples were mixed in a 1:1 ratio and aliquots and cell surface CD27expression was monitored on a flow cytometer while continuously increasing the sample temperature to 37°C using an IR lamp. A temperature of 37°C was reached after 7-8 minutes and kept constant while measuring continued for another 7-8 minutes

### Monitoring P2X7-induced pore formation

BMM, peritoneal macrophages and brain microglia were isolated from P2X4ko and WT mice and WT or P2X4ko cells were stained witheFluor^670^. The labeled and unlabeled cells were mixed in a 1:1 ratio, washed and transferred into RPMI medium and aliquots were subjected to real-time P2X7-induced pore formation analyses using 4’,6-Diamidin-2-phenylindol (DAPI, 1.5μM) uptake as readout for. For this, the baseline DAPI signal was measured by flow cytometry for 2 minutes, then ATP was added to the sample mixture at the indicated concentration and measuring continued for 3-4 minutes.

### Monitoring P2X7-induced calcium uptake

BMM or HEK cells stably transfected with expression plasmids for P2X7k 451L, P2X7k 451P, P2X7a 451L or P2X7a 451P were loaded with 2 μM Fluo-4 (Invitrogen) for 20 min at 4°C and 10 min at 37°C, washed once with FACS buffer and resuspended in PBS supplemented with 0.9 mM CaCl_2_ and 0.49 mM MgCl_2_ (Invitrogen) and analyzed by flow cytometry. An infrared lamp was used to maintain a constant sample temperature of 37°C. The baseline Fluo4 signal was measured for 2 minutes, then ATP was added to the sample at the indicated concentration and measuring continued for 3-4 minutes.

### In vitro migration assay

Th were isolated from WT and P2X4ko mice by FACS and one WT or P2X4ko Th were stained witheFluor^670^. Labeled and unlabeled cells were mixed in a 1:1 ratio, washed and resuspended in RPMI complete medium resuspendiert. 2 x 10^5^ cells in 100 μl were transferred into the upper chamber of a transwell plate (5μm pores, Corrning). The lower chamber was prepared with either 150μl RPMI complete or 150μl RPMI complete containing SDF1α (100ng/ml, Biolegend). The transwell plate was placed in a 37 °C incubator and cells were allowed to migrate for 2 h. Afterwards, cells in the upper and lower chamber were counted.

### Mast cell degranulation assay

Cells were harvested from the peritoneum by lavage with 5 ml cold PBS + EDTA (2 mM, Gibco). After washing, cells were resuspended in RPMI complete medium and aliquots were stimulated with ATP (50 – 800μM) for 10 min at 37°C. Mast cells were identified by flow cytometry as CD11b-FcεR1+ and ATP-induced degranulation of mast cells was evaluated by CD107a externalization.

### Cytokine secretion assay

For in vitro cytokine secretion assays CTL were isolated by FACS from spleen single cell suspensions. 5 x 10^4^ cells were directly sorted in 200 μl RPMI complete medium and stimulated for 24 hours in the presence of phorbol 12-myristate 13-acetate (PMA, 20ng/ml, Invivogen) and ionomycin (1μg/ml, Invivogen). Supernatants were analyzed for 13 cytokines (IFN-γ, TNF-α, IL-2, IL-4, IL-21, IL-22, IL-17A, IL-17F, IL-10, IL-9, IL-5 and IL-13) using the LEGENDplex mouse Th cytokine 13-plex (Biolegend) kit.

### In silico research and statistics

mRNA sequencing data from CD4 T cells of inbred mouse strains was obtained from the immgen database (GSE60337) (Mostafavi et al., 2014). Details of *P2rx7* neighboring genes were obtained from BioMart on ensembl.org (Yates et al., 2020). For statistical analyses, GraphPad Prism 8 was used and two groups were compared using the student’s t test.

## Results

### P2X7 expression levels on T cells in laboratory mouse strains are associated with the P2X7 451P/L polymorphism

The well characterized single nucleotide polymorphism (SNP) rs48804829 leads to a proline (451P) to leucin (451L) exchange at amino acid position 451 in the P2X7 protein. B6 mice express the P2X7^451L^ variant whereas 129 and Balb/c mice express the P2X7^451P^ variant. This SNP affects P2X7 pore formation (Adriouch et al., 2002; Sorge et al., 2012). In order to determine whether this SNP affects P2X7 expression levels, we analyzed the RNA sequencing dataset provided by Mostafavi et al. to compare *P2rx7* mRNA expression levels of CD4^+^ T cells from 23 different mouse strains (Mostafavi et al., 2014). The results show that 451P strains express higher *P2rx7* levels in CD4^+^ T cells, compared to 451L strains (**Fig.1A**). We next analyzed 129, Balb/c, B6 and FVB/N T cells for their P2X7 expression using the P2X7-specific monoclonal antibody Hano44 (Adriouch et al., 2005)and flow cytometry. In agreement with the RNA expression analysis, the results show higher cell surface levels of P2X7 on CD8^+^ cytotoxic T lymphocytes (CTL) and CD4^+^ helper T cells (Th) from 129 and Balb/c mice than from B6 and FVB/N mice (**Fig.1B**). Of note, regulatory T cells (Treg) display high levels of P2X7 on the cell surface in all analyzed mouse strains. Comparison of *P2rx7* mRNA levels from CTL and Th cells of B6 and Balb/c mice confirmed that both Balb/c cell population expressed about 5-fold higher *P2rx7* mRNA levels (**Fig.1C**). Hence, our data shows that the 451P/L *P2rx7* polymorphism is associated with higher/lower P2X7 expression in two major T cell populations.

**Figure 1:**
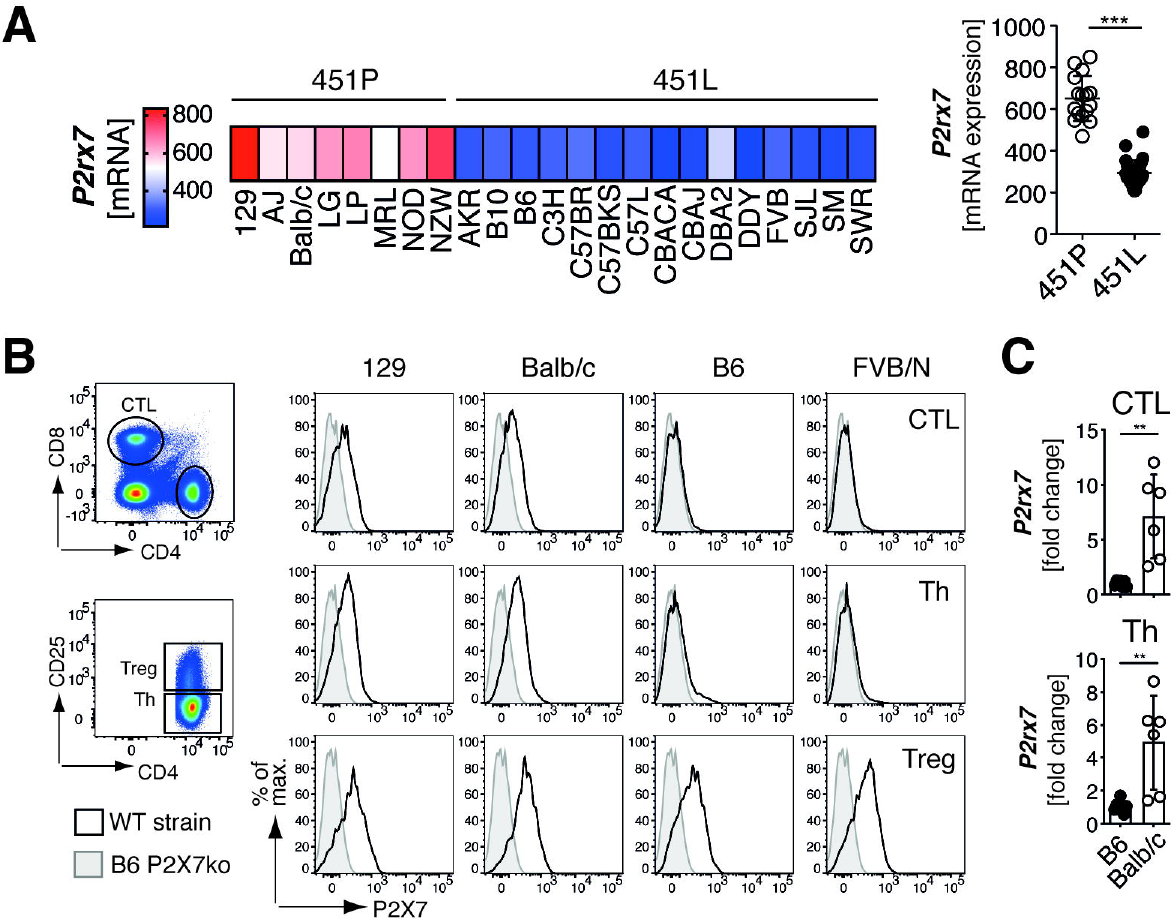
P2X7 expression levels in T cells are associated with the 451P/L polymorphism. **(A)** Comparison of *P2rx7* mRNA expression on CD4^+^ T cells from individual mouse strains expression P2X7 451P (white) or 451L (black). *P2rx7* mRNA data was pool among 451L and 451P strains and compared. mRNA expression data was obtained from www.imgen.org (Heng et al., 2008; Mostafavi et al., 2014). **(B)** Flow cytometric analyses of cell surface P2X7 expression on CD8^+^ cytotoxic T cells (CTL) CD4^+^CD25^-^ helper T cells (Th) and CD4^+^CD25^+^ regulatory T cells (Treg) of 129, Balb/c, B6 and FVB/N mice. **(C)** *P2rx7* mRNA expression was analyzed in CTL and Th from B6 and Balb/c mice (n = 5-6). Statistical comparison of two groups was performed by using the student’s t test (p < 0.05 = * / p < 0.01 = ** / p < 0.001 = ***, ns = no significant).

### P2rx4^tm1Rass^ mice carry the 129-originating P2rx7 gene as passenger mutation

One of the first P2X4-deficient mouse lines, P2rx4^tm1Rass^, was generated by the Rassendren lab in 2006 (Sim et al., 2006). Here, *P2rx4* was targeted in ESCs of the 129 mouse strain and the generated P2X4ko mice were then backcrossed onto the B6 background. Since *P2rx4* and *P2rx7* are adjacent genes on mouse chromosome 5, we hypothesized that the 129-derived P2X7^451P^ is still present in B6-P2X4ko mice (**Fig.2A**). We confirmed this hypothesis by sequencing an amplified fragment flanking rs48804829 (**Fig.2B**). B6-WT and B6-P2X4ko thus differ in both, *P2rx4* and *P2rx7*. In order to obtain mice that differ only in *P2rx4* but not *P2rx7*, we backcrossed P2X4ko mice onto the Balb/c background for 13 generations. This way, we obtained Balb/c-WT and Balb/c-P2X4ko mice which both express P2X7^451P^ in addition to B6-WT and B6-P2X4ko mice that express P2X7^451L^ and P2X7^451P^. We next compared P2X7 expression on T cells of B6 and Balb/c P2X4ko mice and the corresponding WT strains. Here, we observed that CTL and Th but not Treg from B6-P2X4ko mice expressed higher level of P2X7 when compared to B6-WT T cells. In contrast, Balb/c P2X4ko and WT T cells exhibited comparable P2X7 expression levels (**Fig.2C**). Finally, we isolated mRNA from CTL and Th from all four strains and analyzed *P2rx7* mRNA expression. The results show that B6-P2X4ko CTL and Th express 3-6 fold higher level of *P2rx7* mRNA compared to B6-WT T cells. There was no statistically significant difference in *P2rx7* mRNA expression in Balb/c P2X4ko and WT CTL and Th (**Fig.2D**). Of note, frequencies of T cells in blood, lymph nodes and spleen were similar in B6-WT and P2X4ko mice and Balb/c-WT and Balb/c-P2X4ko mice, with a slight but significantly reduced frequency of CTLs in lymph nodes and spleen of B6-P2X4ko mice (**SFig.1**). Taken together, B6-P2X4ko mice carry the P2X7^451P^ variant as passenger mutation, which is associated with a higher P2X7 expression on T cells compared to corresponding WT cells.

**Figure 2:**
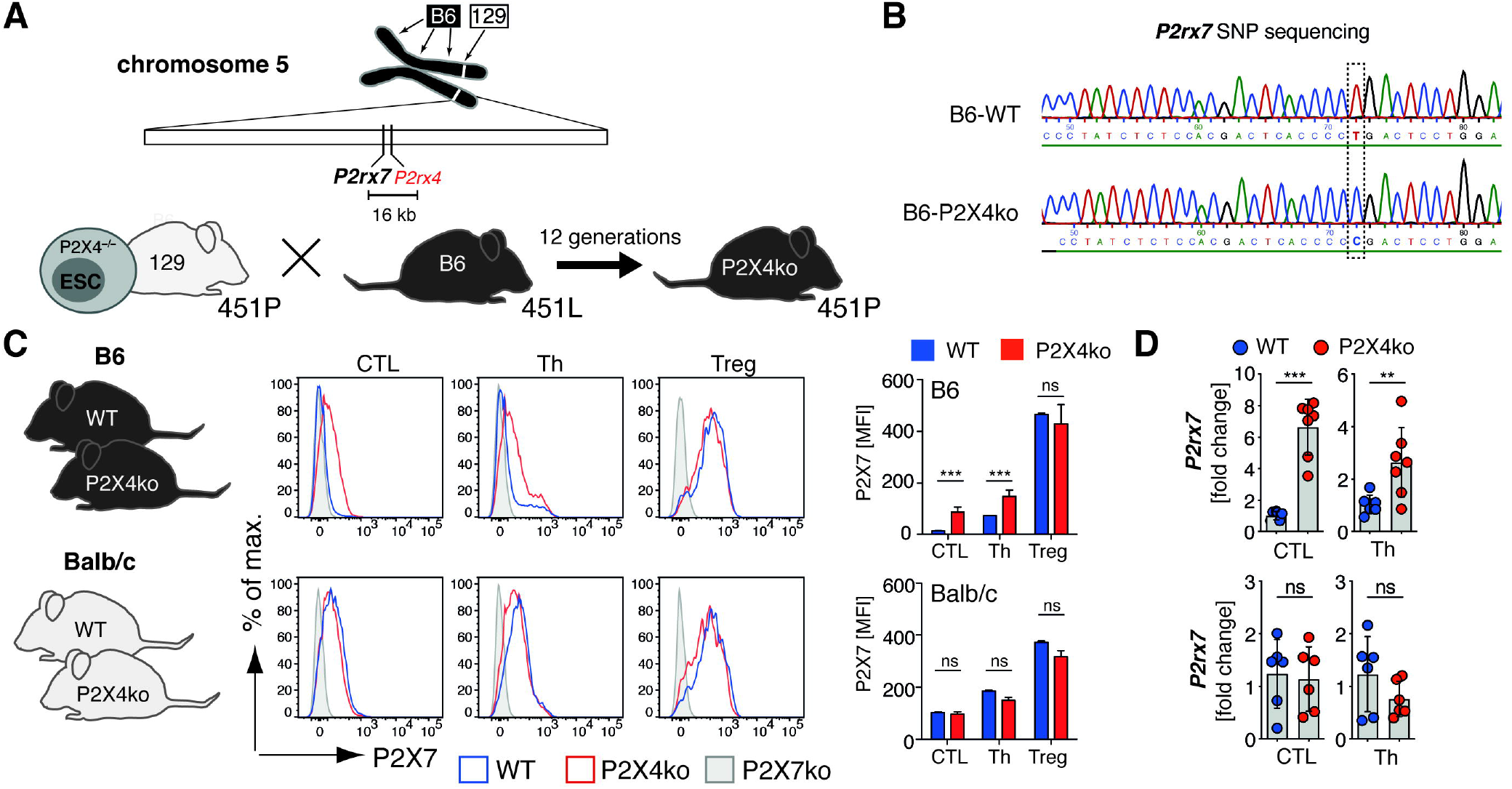
P2X4ko mice harbor a *P2rx7* passenger mutation. **(A)** *P2rx7* and *P2rx4* are neighboring genes on mouse chromosome 5. A non-synonymous SNP in the *P2rx7* gene introduces a single amino acid change in the P2X7 protein: 129 mice express a proline at position 451 (451P) whereas B6 mice express a leucin at position 451 (451L). P2X4ko mice were generated by deleting exon 1 of *P2rx4* in embryonic stem cells derived from a 129 mouse strain. These mice express the 451P variant of P2X7. Backcrossing of P2X4ko onto the B6 background leads to the generation of B6-P2X4ko mice that express the 451P variant, due to a high degree of genetic linkage. **(B)** Sequencing of cDNA obtain from isolated immune cell mRNA confirmed the 451P passenger mutation in the B6-P2X4ko mice. **(C)** Flow cytometric analyses of cell surface P2X7 expression on CTL, Th and Treg of WT and P2X4ko mice on the B6 and Balb/c background. The mean fluorescence intensity (MFI) of P2X7 on the different T cell populations from WT and P2X4ko mice (n = 3) was compared. **(D)** *P2rx7* mRNA of FACS sorted WT and P2X4ko CTL, Th and Treg (B6 and Balb/c) was analyzed (n = 6-7). Statistical comparison of two groups was performed by using the student’s t test (p < 0.05 = * / p < 0.01 = ** / p < 0.001 = ***, ns = no significant).

### B6-P2X4ko T cells are more sensitive to ATP and NAD^+^ compared to B6-WT T cells

P2X7 expression levels likely influence the cellular response to adenosine triphosphate (ATP). To analyze this, we generated stably transfected HEK cell lines that express P2X7^451L^ or P2X7^451P^, each in the two major splice variants (the canonical P2X7a and P2X7k) (Nicke et al., 2009; Schwarz et al., 2012). We first compared ATP-induced calcium responses of these variants by real-time flow cytometry. P2X7k^451L^ HEK cells were labeled with eFLuor^450^ and mixed with unlabeled P2X7k^451P^ HEK cells. Mixed cells were loaded with Fluo4 and were analyzed in parallel. In order to adjust for cell surface levels of P2X7, cells were stained with AF647-conjugated Hano44 and the analysis gate was adjusted on the basis of P2X7 mean fluorescence intensity (MFI). All cell lines responded with a comparably strong and ATP dose-dependent Ca^2+^ influx (**SFig.2A**). Shifting the analysis gates in P2X7^451L^ HEK cells towards a higher P2X7 MFI and of P2X7^451P^ HEK cells towards a lower P2X7 MFI, resulted in increased and decreased calcium signals, respectively (**SFig.2B**). This suggests that the level of cell surface P2X7 influences ATP induced calcium signals and likely also downstream functions. Since CTL and Th cells from B6-P2X4ko mice exhibit higher P2X7 expression levels, it is likely that they also react more sensitive to stimulation with ATP. To test this hypothesis, we incubated splenocytes from B6-WT and B6-P2X4ko mice with rising concentrations of ATP for 10 min at 37°C and measured the loss of CD27 from the cell surface, a well-known readout for P2X7 activation on T cells (Moon et al., 2006; Rissiek et al., 2018a). Strikingly, the majority of B6-P2X4ko CTL and Th responded already to incubation at 37°C with a massive loss of CD27 from their cell surface, in contrast to their B6-WT counterparts (**Fig.3A**). In contrast, Treg cells from B6-WT and B6-P2X4ko mice both responded with a strong loss of CD27, when incubated at 37°C. Likewise, B6-P2X4ko CTL and Th were more sensitive to addition of exogenous ATP, with an ED_50_ of 120μM (P2X4ko) and 225μM (WT) for CTL and 100μM(P2X4ko) and 170μM (WT) for Th cells. Of note, a similar experiment with Balb/c P2X4ko and Balb/c WT splenocytes showed no significant difference in ATP sensitivity between P2X4ko and WT T cells.

**Figure 3:**
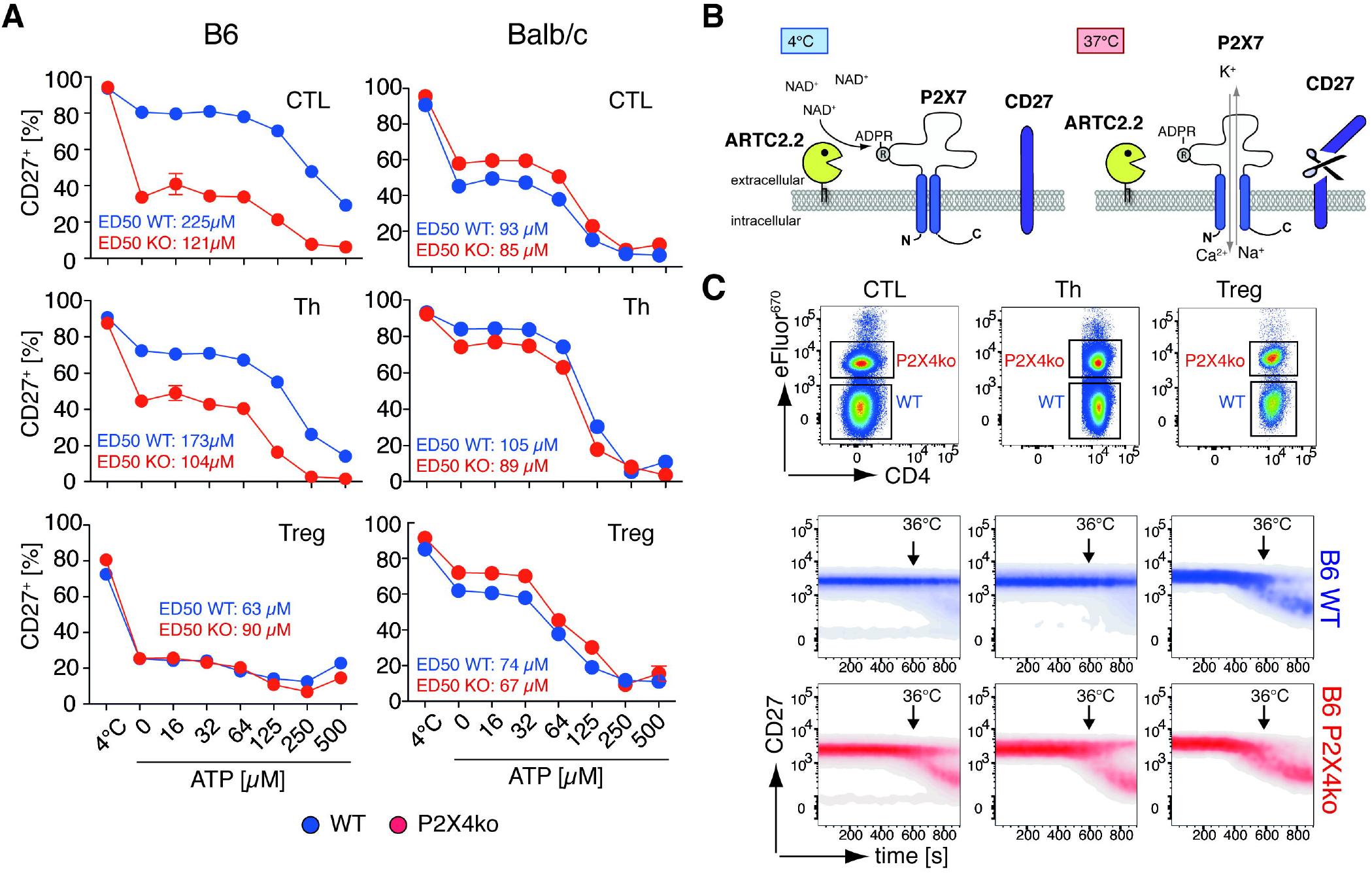
B6 P2X4ko T cells show an increased response to ATP and NAD^+^-mediated P2X7 activation when compared to B6 WT T cells. **(A)** CTL, Th and Tregs from WT (blue) and P2X4ko mice (red) on the B6 and Balb/c background were incubated at 37°C in the presence of 0-500 μM ATP or left at 4°C. CD27 shedding was assessed as surrogate readout for P2X7 activation. ED50 for ATP stimulation was calculated. **(B)** Ecto-ADP-ribosyltransferase ARTC2 can ADP-ribosylate P2X7 at arginine (R) 125. Extracellular NAD^+^, which is release during cell preparation, can serve as substrate for ARTC2.2, even if cells are prepared at 4°C. Bringing the cells back to 37°C induces gating of P2X7 inducing shedding of CD27. **(C)** B6 P2X4ko splenocytes were labeled with eFluor^670^ and mixed with unlabeled B6 WT splenocytes. Mixed cells were stained with anti-CD27 and cell surface CD27 was monitored over time on CTL, Th and Treg while increasing the temperature up to 36°C.

The loss of CD27 during incubation at 37°C is most likely due to activation of P2X7 by ADP-ribosylation in response to extracellular NAD released during cell isolation (**Fig.3B**). This effect has been well described for Treg, natural killer T cells (NKTs), tissue-resident memory T cells (Trm) and T follicular helper cells (Georgiev et al., 2018; Rissiek et al., 2018b; 2014; Scheuplein et al., 2009) which co-express P2X7 and ARTC2.2 (Glowacki et al., 2002). While ADP-ribosylation usually occurs at 4°C, ADP-ribosylation-induced gating of P2X7, however, needs temperatures above 30°C (Scheuplein et al., 2009). To better visualize the temperature dependence of this process, we labeled B6-P2X4ko splenocytes with eFluor^670^, mixed them with unlabeled B6-WT splenocytes at 4°C, and then monitored the loss of CD27 on CTL, Th and Treg cells at elevating temperatures using real-time flow cytometry. During the measurement, the sample was brought to 36°C while the CD27 signal was continuously monitored. Once the cells reached temperatures close to 36°C, the CD27 signal decreased in a fraction of cells (**Fig.3C**). Apparently, this loss of CD27 was more prominent in CTL and Th from B6-P2X4ko compared to corresponding B6-WT cells (**Fig.3C**). Again, Tregs from both, B6-P2X4ko and B6-WT mice, reacted with a pronounced loss of CD27 in this experimental setup. Of note, ARTC2.2 expression levels were comparable among individual T cell populations from P2X4ko and corresponding WT mice (**SFig.3**), as determined by flow cytometry.

### The increased sensitivity of B6-P2X4ko T cells towards NICD confounds results from functional assays

The ultimate consequence of ADP-ribosylation-mediated P2X7 activation is NICD (Seman et al., 2003). Cell preparation related ADP-ribosylation of P2X7 on T cells can be prevented by injecting the ARTC2.2 blocking nanobody s+16a prior to sacrificing the mice (**Fig.4A**) (Koch-Nolte et al., 2007, Rissiek et al., 2014). Since CTL and Th from B6-P2X4ko mice are sensitive to P2X7 induced CD27 shedding at 37°C, we next tested whether this also results in an increased susceptibility towards NICD of FACS-sorted CTL and Th from mice that had been treated or not with s+16a. As a readout for cell death, we measured staining of cells with propidium iodide (PI). We observed that only a minor fraction of B6-WT CTL and Th (20-30%) underwent NICD. In contrast, 40-60% of B6-P2X4ko CTL and Th were PI^+^ after 2h of incubation at 37°C. Of note, injection of s+16a prevented NICD of B6-WT and B6-P2X4ko CT and Th (Fig.4A). Next, we compared cytokine production and migration of CTL and Th from mice that had been treated or not with s+16a. For cytokine analyses, we stimulated FACS-sorted CTL with phorbol-12-myristat-13-acetat (PMA) / ionomycin. When comparing CTL from untreated B6-P2X4ko and B6-WT mice, it appeared that P2X4-deficiency dampened secretion of IL-2, IFNγ and TNFα. However, when ADP-ribosylation during cell preparation was prevented by s+16a injection, CTL from both mouse lines produced comparable amounts of all three cytokines (**Fig.4B**). Since P2X4 has been attributed a role in T cell migration (Ledderose et al., 2018), we next compared B6 WT and P2X4ko Th in an *in vitro* migration assay using a trans-well chamber and SDF1α (CXCL12) as chemoattractant in the bottom chamber. The presence of SDF1α markedly increased the migration of Th cells from the top to the bottom wells, however, B6-P2X4ko Th cells seemed to migrate less compared to B6-WT Th (**Fig.4C**). This difference in migration was not observed with cells from mice that had been treated with the ARTC2.2 blocking nanobody s+16a. This suggests that increased NICD susceptibility rather than a P2X4-mediated effect on migratory capacity causes the difference in Th migration. This conclusion is supported by the finding that WT and P2X4ko Th cells from Balb/c mice, which do not differ in NICD susceptibility, also did not differ in their migration capacity towards SDF1α (Fig.4C). Finally, we performed *in vivo* adoptive transfer experiments where we mixed differentially labeled B6-WT and B6-P2X4ko CTL harvested from mice that had been treated or not with s+16a in a 1:1 ratio and transferred them into RAG1ko mice. 24h after adoptive transfer, spleens of the injected RAG1ko mice were analyzed for the presence of the injected CTLs. Here, we retrieved about 2-fold more B6-WT than B6-P2X4ko CTL from the RAG1ko spleens, again suggesting a deficit in migration capacity of the B6-P2X4ko CTL (**Fig.4D**). However, when B6-WT and B6-P2X4ko CTL were harvested from s+16a treated mice, B6-WT and B6-P2X4ko CTL were retrieved at a ratio of almost 1:1.

**Figure 4:**
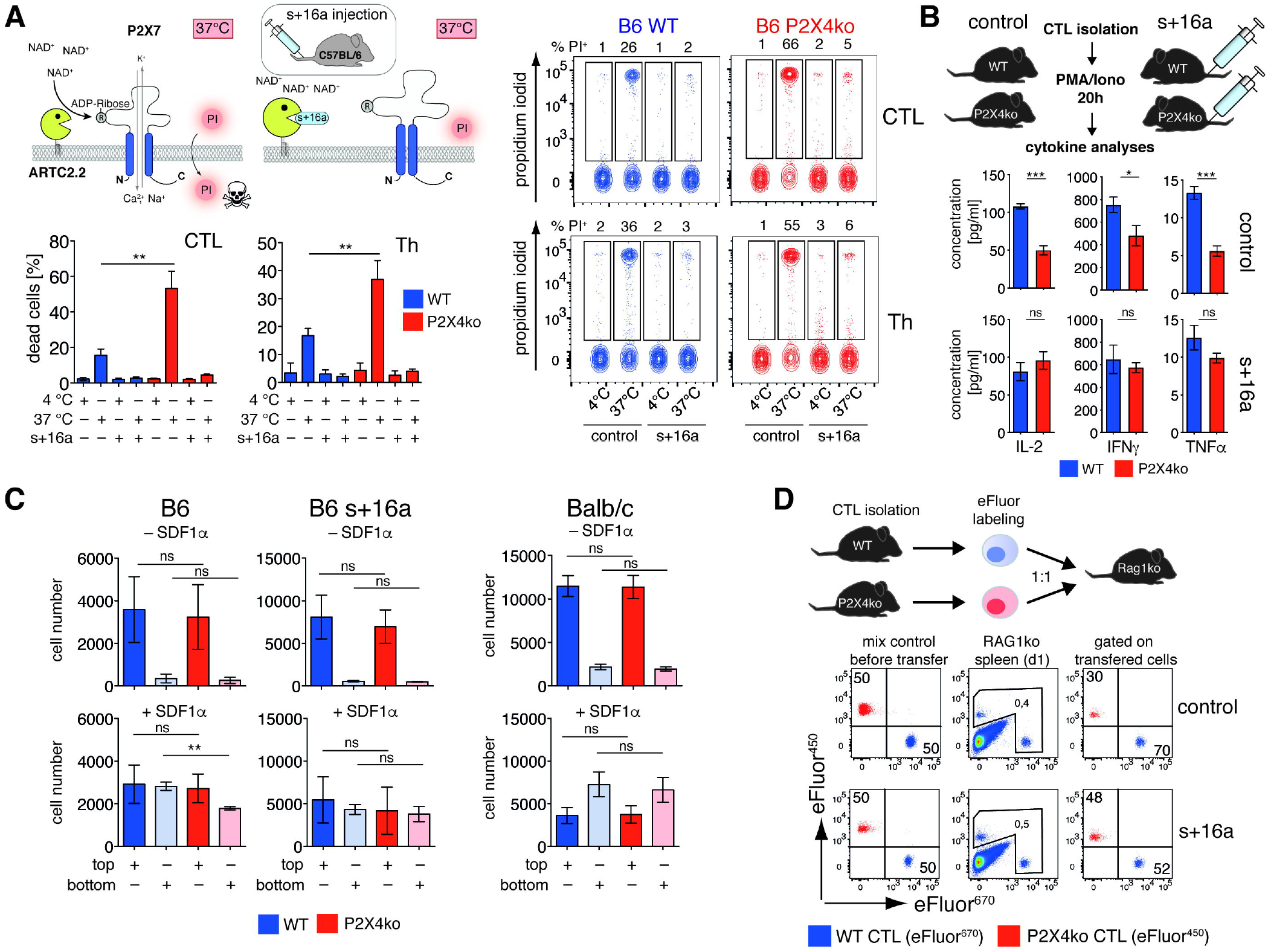
The enhanced NICD of B6-P2X4ko T cells can lead to misinterpretation of functional assays. **(A)** B6 WT and B6 P2X4ko mice were injected or not with anti-ARTC2.2 nanobody s+16a in order to prevent P2X7 ADP-ribosylation during cell preparation. CTL and Th from spleen were FACS sorted and obtained cells were incubated at 37°C for 2h or left at 4°C. Cell vitality was measured by propidium iodide (PI) uptake (n = 3). **(B)** B6 WT and P2X4ko mice were injected (s+16a) or not (control) with anti-ARTC2.2 nanobody s+16a. 5 x 10^4^ CTL were FACS sorted and stimulated *ex vivo* with PMA/ionomycin for 20h. IL-2, IFNγ and TNFα in the supernatants were measured by cytometric bead array. **(C)** 1×10^5^ Th from P2X4ko mice (B6, B6 treated with s+16a or Balb/c) and 1×10^5^ eFluor^670^ labeled Th from corresponding WT mice were mixed and transferred to the top well of a trans-well chamber in order to perform a comparative migration assay towards SDF1α, which was used as chemoattractant in the bottom well. Migration of Th into the bottom was determined by counting living cells in the top and bottom well after 2h of incubation at 37°C. **(D)** Comparative adoptive transfer of CTL obtained from B6 WT and P2X4ko mice that have been treated or not with s+16a was performed. Cells were differentially labeled with eFluor^670^ and eFluor^450^ and 4×10^5^ cells mixed in a 1:1 ratio were i.v. injected into RAG1ko mice. Spleens were analyzed after 24h for the presence of transferred mixed cells. Statistical comparison of two groups was performed by using the student’s t test (p < 0.05 = * / p < 0.01 = ** / p < 0.001 = ***, ns = no significant).

The results with these three functional assays demonstrate that increased P2X7 expression rather than loss of P2X4 alter several functions of Th and CTL on the B6 background.

### P2X7 expression and function of innate immune cells from P2rx4^tm1Rass^ are downmodulated independently of the 129-derived P2rx7 passenger mutation

Since the *P2rx7* passenger mutation in B6-P2rx4^tm1Rass^ mice influences P2X7 expression levels in T cells, we further analyzed the impact of the *P2rx7* passenger mutation on innate immune cells known to express P2X7. For this, we analyzed P2X7 expression on peritoneal macrophages and mast cells, spleen macrophages and brain microglia obtained from B6-WT and B6-P2X4ko mice. Since sex-related differences have been described for P2X7 expression by innate immune cells (Crain et al., 2009; Guneykaya et al., 2018), we separately analyzed cells from male and female mice. The results of flow cytometry analyses reveal that peritoneal macrophages and mast cells as well as splenic macrophages from B6-P2X4ko mice express much less P2X7 on their cell surface compared to corresponding cells from B6-WT mice (**Fig.5A**). This observation was made in both, male and female mice. In contrast, P2X7 expression was comparable among B6-WT and B6-P2X4ko microglia. Strikingly, when comparing Balb/c-WT and Balb/c P2X4ko, reduced P2X7 expression was observed in P2X4ko macrophages, mast cells, and microglia (Fig.5A). Since bone marrow-derived innate immune cells are widely used in immunological research we generated bone marrow derived dendritic cells (BMDC), bone marrow derived macrophages (BMM) and bone marrow derived mast cells (BMMC) derived from B6 and Balb/c WT and P2X4ko mice and analyzed their P2X7 expression. Again, B6 and Balb/c P2X4ko cells exhibited lower P2X7 expression levels compared to the corresponding WT cells (**SFig.4**). Similar results were obtained with intracellular staining of P2X7 in peritoneal macrophages and mast cells (**Fig.5B**). Consistently, *P2rx7* mRNA expression analyses revealed lower *P2rx7* mRNA levels in BMM, BMDC, BMMC and peritoneal macrophages from P2X4ko mice of both strains when compared to the corresponding WT animals (**Fig.5C**). Of note, microglia mRNA expression analyses reflected the results from flow cytemtry experiments. These findings likely are a consequence of the strategy used to manipulate the gene locus rather than related to the introduced *P2rx7* passenger mutation. We further checked the expression level of other *P2rx7* neighboring genes (*Camkk2, ATP2a2, Kdm2b, Anapc5* and *Anapc7*) in B6 and Balb/c P2X4ko mice. However, apart from a slight but significant reduction of *Camkk2* expression in B6-P2X4ko BMDC in comparison to B6-WT BMDC, there were no significant differences in expression of other *P2rx7* neighboring genes in all analyzed cell populations (**SFig.5**). We next checked the functional impact of the reduced P2X7 expression level in B6 and Balb/c P2X4ko innate immune cells. Direct flow cytometric comparison of P2X4ko and WT BMM, distinguishable by eFluor^670^ labeling, towards ATP-evoked calcium influx revealed that B6 and Balb/c P2X4ko BMM show slower uptake of calcium after ATP stimulation (**Fig.5D**). Further, B6 and Balb/c P2X4ko and WT BMM, peritoneal macrophages and brain microglia were analyzed for ATP induced pore formation. The results resemble the ones from the P2X7 expression analyses: B6 and Balb/c P2X4ko BMM and peritoneal macrophages show a slower uptake of DAPI in response to ATP stimulation. Microglia from B6-P2X4ko mice did not differ in ATP-induced pore formation compared to B6-WT, whereas Balb/c-P2X4ko microglia exhibit a reduced DAPI uptake compared to Balb/c-WT microglia. For peritoneal mast cells, we evaluated whether reduced P2X7 levels on the cell surface affected ATP induced degranulation, as evidenced by ATP-induced externalization of CD107a (Lamp1). Comparative dose response analyses of B6 and Balb/c P2X4ko and WT mast cells revealed that P2X4ko mast cells from both strains need twice the amount of ATP in order to induce a comparable degree of CD107a externalization as corresponding WT mast cells (**Fig.5E**).

**Figure 5:**
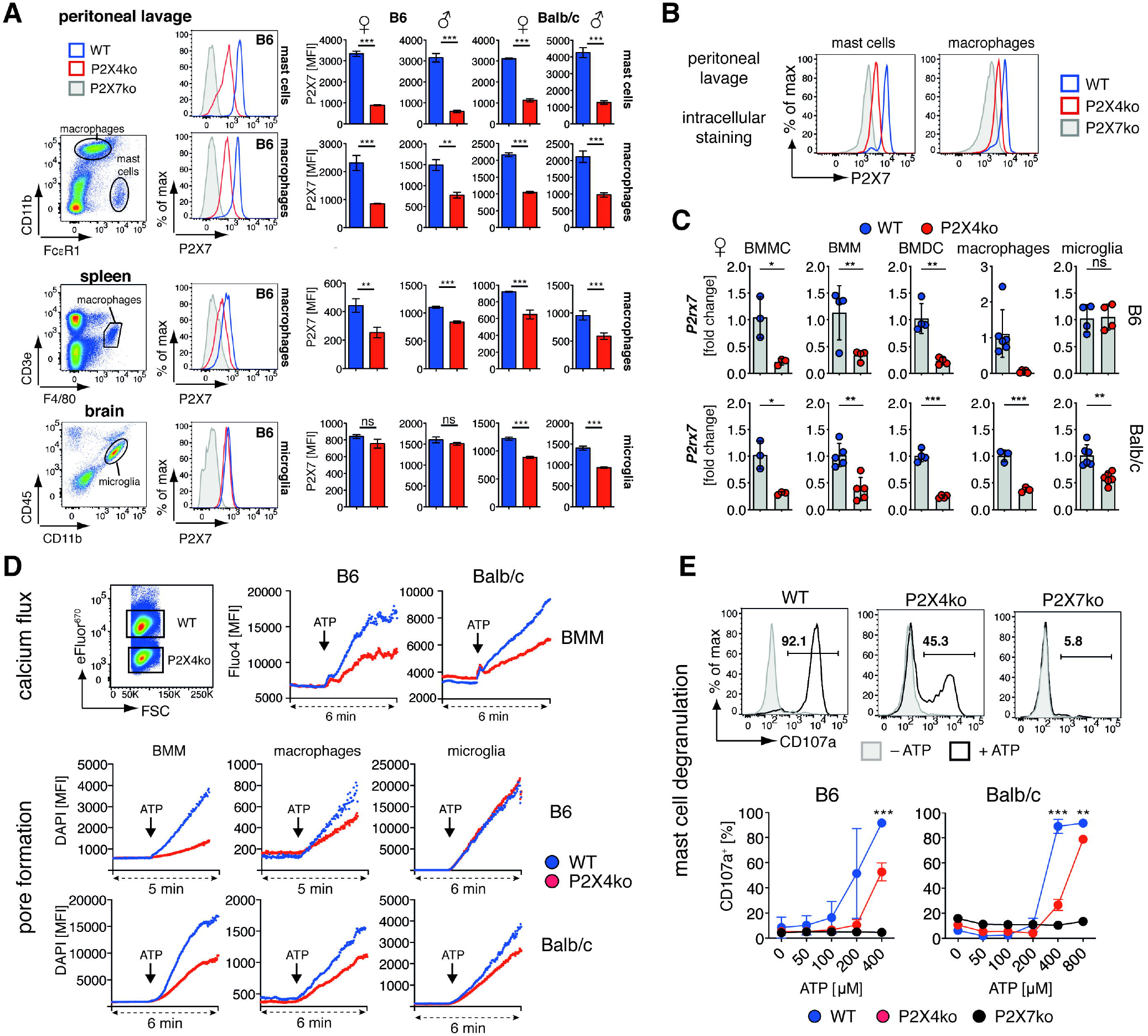
P2X4ko innate cells show reduced P2X7 expression and function. **(A)** P2X7 expression was monitored by flow cytometry on peritoneal macrophages (CD11b^+^Fcγr1^-^), peritoneal mast cells (CD11b^-^Fcγr1^+^), spleen macrophages (CD3e^-^F4/80^+^), and brain microglia (CD11b^+^CD45^int^) from female and male WT and P2X4ko mice on the B6 and Balb/c background (n = 3). **(B)** Intracellular P2X7 expression was measured in peritoneal mast cells and macrophages from B6 WT and P2X4ko mice. **(C)** BMMC, BMM, BMDC, peritoneal macrophages and microglia mRNA obtained from female B6 WT and P2X4ko or Balb/c WT and P2X4ko mice was analyzed for *P2rx7* expression. **D)** Calcium influx was monitored in B6 and Balb/c WT and P2X4ko BMM. Cells were loaded with Fluo4 and intracellular calcium was monitored via real-time flow cytometry (2 min baseline / 4 min + 1.5 mM ATP). Pore formation was monitored in B6 and Balb/c WT and P2X4ko BMM, peritoneal macrophages and brain microglia. DAPI uptake was measured by real-time flow cytometry (2 min baseline / 3-4 min + 1.5 mM ATP). **(E)** Degranulation in response to ATP stimulation was measured in B6 and Balb/c WT and P2X4ko peritoneal mast cells by analyzing cell surface CD107a presence. Statistical comparison of two groups was performed by using the student’s t test (p < 0.05 = * / p < 0.01 = ** / p < 0.001 = ***, ns = no significant).

In summary, P2X7 expression and, therefore, the sensitivity towards ATP stimulation was reduced in all analyzed P2X4ko innate immune cell populations except for microglia on the B6 background, when compared to corresponding WT cells. This effect likely is independent of the *P2rx7* passenger mutation since it was observed in B6 and Balb/c P2X4ko mice.

### Other transgenic mice also carry the P2rx7 passenger mutation

A genetic distance of 2 Mb in the mouse genome is, on average, equivalent to 1 centiMorgan (cM) (Bryant et al., 2018). This implicates for 129-dervived transgenic mice that genes in a 2 Mb maximum distance of the targeted gene are retained with a theoretical probability of 90% after 10 generations of backcrossing. *P2rx4* is with a distance of only 16 kb the nearest neighbor gene of *P2rx7*. However, there are 94 other characterized, proteincoding genes within 2 megabases (Mb) upstream and downstream of *P2rx7*. A search in the Mouse Genome Informatics (MGI) database resulted in 96 mouse strains generated from 129-derived ESCs in which genes within +/− 2MB of *P2rx7* had been targeted (**Tab.1**). To test our hypothesis that the *P2rx7* passenger mutation will affect T cell functions in mice of neighboring genes, we analyzed nine other mouse strains with targeted *P2rx7* neighboring genes on the B6 background for the presence of the 129-derived *P2rx7* passenger mutation. Results of the SNP rs48804829 sequencing revealed that eight of the nine analyzed strains harbor the 129-derived *P2rx7* passenger mutation (**Fig. 6A**). Only in the Atxn2^tm1.1Aub^ mouse strain the 129-derived P2X7^451P^ was exchanged by the B6-derived P2X7^451L^. In order to reproduce the phenotypical and functional consequences of the *P2rx7* passenger mutation described for P2rx4^tm1Rass^, we evaluated T cells from B6-Hvcn1^Gt(RRRN293)Byg^ (B6-Hvcn1ko) mice for the expression of P2X7. Here, we obtained a similar result as for *P2rx4^tm1Rass^*: the P2X7 cell surface expression was significantly higher on B6-Hvcn1ko CTL and Th compared to B6-WT (**Fig.6B**). Similarly, *P2rx7* mRNA levels were increased in B6-Hvcn1ko CTL and Th in comparison to B6-WT CTL and Th (**Fig.6C**). As a consequence, B6-Hvcn1ko CTL and Th responded more pronounced to preparation related P2X7 ADP-ribosylation, as indicated by a massive shedding of CD27 upon incubation at 37°C, when compared to B6-WT T cells (**Fig.6D**). Since innate immune cell populations from P2rx4^tm1Rass^, especially peritoneal macrophages and mast cells, exhibited much lower level of P2X7 expression compared to corresponding B6 WT cells, we analyzed peritoneal macrophages and mast cells from B6-Hvcn1ko mice for their P2X7 expression level. We observed a similar level of P2X7 on macrophages and only a slight but significant reduction of P2X7 protein expression on mast cells. The absence of a pronounced dysregulation of P2X7 expression in Hvcn1ko innate immune cells further suggests that the observed alterations in P2X7 expression in these cells in P2rx4^tm1Rass^ are due to gene locus modifications, e.g. the inactivated *P2rx4* gene, introduced during gene targeting.

**Figure 6:**
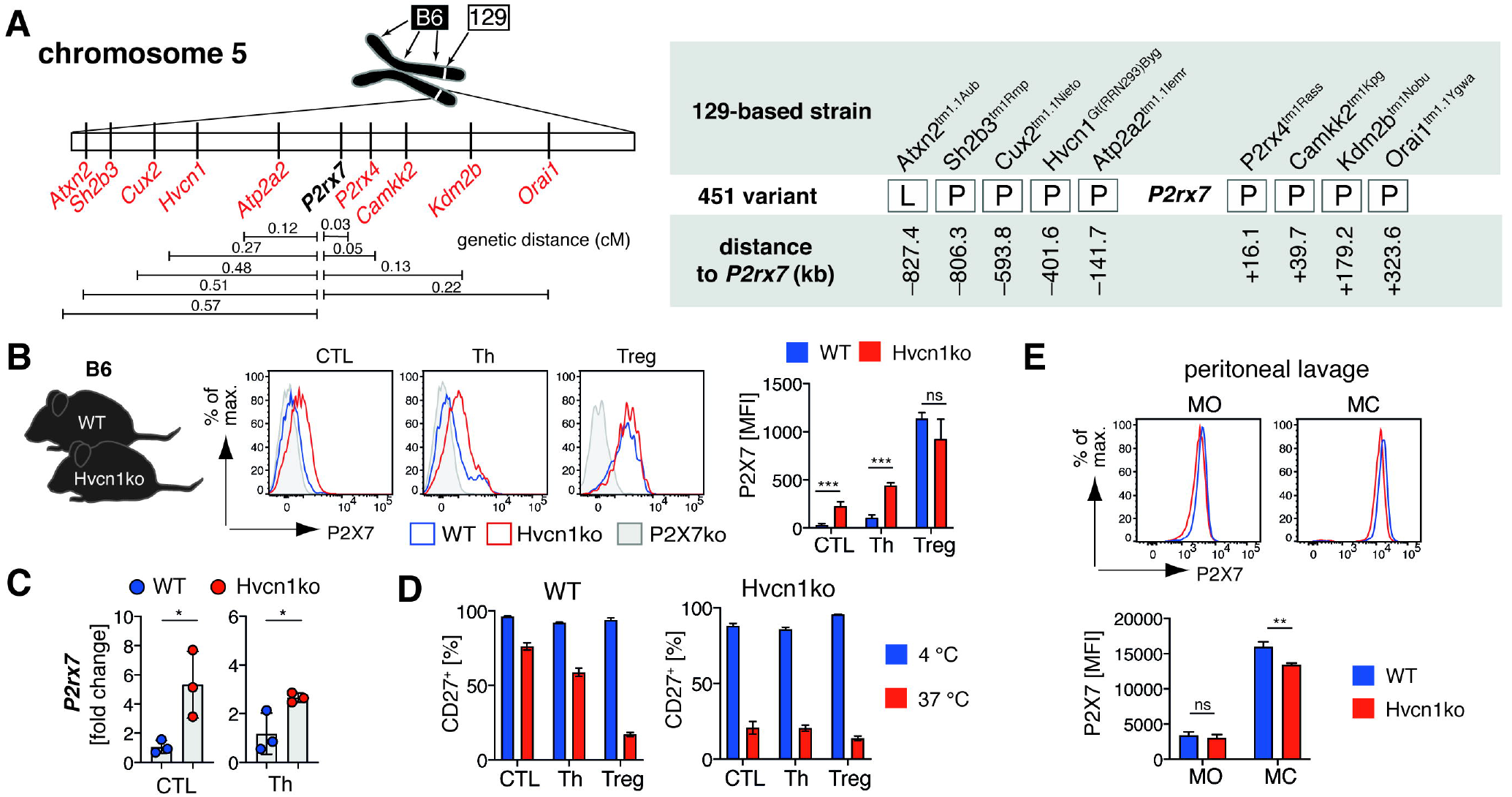
The 129-derived *P2rx7* passenger mutation is present in other transgenic mouse strains. **(A)** 9 gene targeted mouse strains with targeted genes in close proximity of *P2rx7* were analyzed for the presence of the SNP leading to the 451L/P polymorphism. **(B)** P2X7 expression was determined on CTL, Th and Treg from B6-Hvcn1ko carrying the 129-derived P2rx7 passenger mutation (n = 3). **(C)** *P2rx7* mRNA expression was analyzed in CTL and Th from B6-Hvcn1ko and B6-WT mice cells. **(D)** CTL, Th and Treg from Hvcn1ko and B6-WT were incubated at 4°C and 37°C for 15 min in order to measure the impact of P2X7 ADP-ribosylation by shedding of CD27. **(E)** Peritoneal macrophages (MO) and mast cells (MC) from B6-Hvcn1ko and B6-WT mice were analyzed for P2X7 expression (n = 3). Statistical comparison of two groups was performed by using the student’s t test (p < 0.05 = * / p < 0.01 = ** / p < 0.001 = ***, ns = no significant).

## Discussion

Passenger mutations are a phenomenon occurring in genetically modified mice when the genetic background of the used ES cells and the genetic background of the line in which the modified gene was backcrossed are not identical. Even after backcrossing for more than 12 generations, the flanking regions of the modified gene often contain multiple foreign genes of ES cell origin, some of which can be differentially expressed between the transgenic strain and the supposed WT strain. Further, functionally inactive 129-derived passenger genes can introduce an experimental bias, as in the example of the Caspase-1 knockout mouse (Casp1^tm1Flvw^). Here, the 129-derived ES cells used to generate Casp1^tm1Flvw^ (Kuida et al., 1995) introduced a defective *Casp11* gene in close proximity of the knocked out *Casp1* gene, thereby transferring resistance to otherwise lethal LPS injection to Casp1^tm1F1vw^ mice. The role of the inactive *Casp11* passenger gene in LPS resistance was unnoticed for more than 15 years until Kayagaki and colleagues revealed this important fact (Kayagaki et al., 2011). The defective *Casp11* gene was also found in other transgenic mice generated with 129-derived ES cells, such as Panx1^tm1.1Vshe^ (Dvoriantchikova et al., 2012), again conferring these transgenic mice resistance to LPS (Berghe et al., 2015). These examples illustrate the importance of screening transgenic mice with a congenic background for passenger mutations in order to avoid experimental bias.

In this study we demonstrate that the expression of P2X7 on conventional helper T cells as well as on cytotoxic T cells differs among laboratory mouse strains. Interestingly, P2X7 expression was high on T cells from strains carrying the 451P variant of P2X7 whereas P2X7 expression was low on T cells from strains carrying the 451L variant resulting from the SNP rs48804829. Regarding T cells, the 451L P2X7 variant of B6 mice is often referred to as less sensitive or loss-of-function variant, compared to the e.g. the Balb/c 451P P2X7 variant (Bartlett et al., 2014; de Campos et al., 2012; Proietti et al., 2014). However, the lower cell surface levels of P2X7 on B6 T cells compared to Balb/c T cells also contribute to the reduced ATP / NAD^+^ sensitivity. This is underlined by our findings with P2X7-transfected HEK cells, which show comparable ATP-induced calcium influx of 451L and 451P variants when adjusted for P2X7 expression levels.

Since T cells from 129 mice (451P) and B6 mice (451L) mice differ in their P2X7 expression level, we hypothesized that this difference would be passed on as passenger mutation to the B6 background from 129-based mice with targeted genes in the *P2rx7* genetic neighborhood. For B6 mice in which the nearest neighbor gene of *P2rx7, P2rx4*, had been targeted in 129 ES cells, we found that these B6 P2X4ko mice 1) carry *P2rx7* 451P as passenger gene, 2) express higher levels of P2X7 on CD4 and CD8 T cells, when compared to B6 WT, 3) are more sensitive to treatment with exogenous ATP and NAD+, and 4) are more prone to ADP-ribosylation of P2X7 during cell preparation resulting in NAD-induced cell death (NICD) in a higher fraction of cells compared to B6 WT. Increased sensitivity of NICD is receiving more and more attention, since important T cell populations that co-express high level of P2X7 and ARTC2.2, such as Tregs, NKTs, Tfh and Trm, are highly affected by ADP-ribosylation during cell preparation (Georgiev et al., 2018; Hubert et al., 2010; Proietti et al., 2014; Rissiek et al., 2018b; 2014; Stark et al., 2018). The analyses and functional characterization of these cell populations has been greatly improved by the use of the ARTC2.2-blocking nanobody s+16a, which prevents ADP-ribosylation of P2X7 during cell preparation and preserves the vitality of the prepared cells in functional assays and during adoptive transfer (Koch-Nolte et al., 2007; Rissiek et al., 2014). Similarly, injection of s+16a into P2X4ko mice preserved the vitality of the NAD-sensitive P2X4ko T cells. Otherwise, the higher loss of P2X4ko T cells during functional assays *in vivo* and *in vitro*, due to NICD, skews experimental results, as demonstrated by our cytokine expression and migration analyses. This likely holds also for previously reported studies with T cells from P2rx4^tm1Rass^ mice, e.g. where polyclonally activated P2X4ko T cells exhibited less proliferation and granzyme B expression compared to T cells from B6 WT mice (Ventre et al., 2017).

We found that eight of nine other transgenic strains on the B6 background derived from 129 ES cells with targeted genes in the *P2rx7* neighborhood carried the 129-dervied *P2rx7* passenger gene. We had further access to one of these strains, B6-Hvcn1^Gt(RRRN293)Byg^ mice, and were able to reproduce the phenotype of P2rx4^tm1Rass^ T cells, suggesting that our findings can be extrapolated to other transgenic mice carrying the *P2rx7* passenger gene. *Orai1*, for example, plays an important role in T cell function (Nohara et al., 2015) and is also a neighboring gene of *P2rx7*. Studies have demonstrated that genetic ablation of *Orai1* in mice on a pure B6 background e.g. Orai1^tm1Rao^ or Orai1^tm1Fesk^ leads to a diminished T cell cytokine production (Kim et al., 2014; McCarl et al., 2010), underlining the importance of *Orai1* in T cell function. In our study, we could identify the 451P *P2rx7* passenger mutation in the 129-derived Orai1^tm1.1Ygwa^ strain. These mice were used in a study to generate T cell specific B6 Orai1ko mice by crossing them with CD4-Cre mice. Interestingly, CD8 T cells from these T cell-specific B6 Orai1ko were less potent in killing peptide loaded EL-4 tumor cells *in vitro* compared to CD8 T cells from B6 WT mice (Kim et al., 2017). Further, T cells from the 129-based Orai1^Gt(XL922)Byg^ mouse also exhibited reduced IL-2 and IFNγ production in response to polyclonal T cell receptor stimulation (Vig et al., 2008). It is conceivable that these mice also harbor the 129-derived *P2rx7* passenger gene and their T cells are consequently also more sensitive to NICD than the WT controls. It would be interesting to determine, whether treatment with the ARTC2.2 blocking nanobody s+16a has any impact on the outcome of these experimental settings.

Further, it is worth noting that when working with conditional knockouts, the control group can have a great impact on the results and interpretation when it comes to the impact of passenger mutations: A comparison of Cre-floxed and Cre+ floxed mice would “silence” the impact of a passenger mutation, since it is present in both mice. In contrast, a comparison of Cre+ floxed and Cre+ WT or pure WT mice can introduce an experimental bias triggered by a passenger mutation. Finally one has to keep in mind that while a transgenic strain in one animal facility can carry a 129-derived passenger gene, the same strain bred in another animal facility might have successfully replaced the passenger mutation during backcrossing.

Since a passenger gene is present in all nucleated cells and P2X7 plays an important role in innate immunity we also analyzed innate immune cells from P2rx4^tm1Rass^ mice towards a potential impact of the *P2rx7* passenger gene. Surprisingly, we observed a much lower expression of *P2rx7* mRNA and lower levels of P2X7 on the cell surface in many but not all analyzed innate cell populations of P2X4ko mice on the B6 and Balb/c genetic background in comparison to their corresponding WT controls. This finding was consistent in both male and female mice. Similar observations have been made in prior studies: BMDCs from P2rx4^tm1Rass^ were reported to exhibit a decreased *P2rx7* mRNA expression (Zech et al., 2016) and macrophages from P2rx4^tm1Rass^ showed a reduced pore formation capacity in response to ATP stimulation (Pérez-Flores et al., 2015). At last, our finding that microglia from B6 P2X4ko mice express similar level of P2X7 compared to B6 WT also confirms an earlier report (Ulmann et al., 2008). This low P2X7 expression in innate cells of P2X4ko mice could be an effect mediated by *P2rx4* deficiency. Alternatively, insertion of the neomycin (neo) selection cassette in the *P2rx4* locus might also expression of the neighboring *P2rx7* gene (Pham et al., 1996; Rijli et al., 1994). This has been demonstrated, for example, in case of *Gzmb* knockout mice: a first generation of *Gzmb*-targeted mice (Gzmb^tm1Ley^) had a retained neo cassette and exhibited reduced expression of the neighboring genes *Gzmc* and *Gzmf*, whereas the second generation of *Gzmb* knockout mice (Gzmb^tm2Ley^) was generated on the basis of the loxP/Cre system and showed normal expression of *Gzmc* and *Gzmf* (Revell et al., 2005). To answer this question in the P2X4 ko mice, one would need to compare P2X7 expression on innate immune cells from the different P2X4-deficient strains. Interestingly, *P2rx7* mRNA expression level in BMMCs from the P2rx4^tm1Ando^ strain (Yamamoto et al., 2005) where comparable to those of B6 WT BMMCs (Yoshida et al., 2020).

In conclusion, our results emphasize the importance of a thorough analysis of the genetic neighborhood of 129-based transgenic mice on the B6 background. As demonstrated for P2rx4^tm1Rass^, a passenger mutation in the neighboring *P2rx7* gene can introduce an experimental bias. Additionally, gene targeting can also affect the expression of neighboring genes as shown for P2rx4^tm1Rass^ innate immune cells. Apart from immune cells, which were the focus of this study, it would be worthwhile to evaluate the impact of the *P2rx7* passenger mutation on other P2X7 expressing cell populations such as astrocytes, neurons or adipocytes.

## Acknowledgments

The authors would like to thank the following colleagues for providing genomic DNA samples of transgenic mice: Georg Auburger (Atxn2^tm1.1Aub^), Satoshi Takaki and Meena S Madhur (Sh2b3^tm1Rmp^), Marta Nieto and Elia Marcos Grañeda (Cux2^tm1.1Nieto^), Carmella Evans-Molina (Atp2a2^tm1.1Iemr^), David Carling (Camkk2^tm1Kpg^), Nobuaki Yoshida and Manabu Ozawa (Kdm2b^tm1Nobu^), and Yousang Gwack (Orai1^tm1.1Ygwa^). The authors would like to thank the HEXT FACS Core Facility (Hamburg, Germany) for cell sorting.

## Conflict of interest

FK-N receives royalties from sales of antibodies developed in the lab via MediGate GmbH, a 100% subsidiary of the University Medical Center, Hamburg. All other authors declare that the research was conducted in the absence of any commercial or financial relationships that could be construed as a potential conflict of interest.

## Authors contributions

BR and ME-L performed all experiments and analyzed and interpreted the data. YD generated P2X7 stably transfected HEK cells. FR, LU, FU and MAF provided essential material. AN, FK-N and TM supervised the experiments and assisted with data interpretation and manuscript preparation. BR assembled the figures and wrote the manuscript, which has been reviewed by all authors.

## Funding

This work was funded by the Deutsche Forschungsgemeinschaft (DFG, German Research Foundation–Project-ID: 335447717—SFB1328, A06 and A16 to MAF, A10 and Z02 to FK-N, A13 to TM, and A15 to AN), the “Hermann und Lilly Schilling-Stiftung fu□r Medizinische Forschung” to TM, and a faculty grant to BR (FFM, NWF-17/07).

---

**Supplementary figure S1: T cell frequencies are comparable among P2X4ko and WT mice.** Frequencies of CTL, Th and Treg (n = 3-5) in relation to all CD3^+^ T cells was determined in blood, peripheral lymph nodes and spleen of B6 and Balb/c P2X4ko (red) and WT mice (blue). Statistical comparison of two groups was performed by using the student’s t test (p < 0.05 = * / p < 0.01 = ** / p < 0.001 = ***, ns = no significant).

**Supplementary figure S2: The intensity of the calcium influx depends on the expression level of P2X7. (A)** HEK cells stably transfected with P2X7 451L or 451P (P2X7k or P2X7a splice variant) were distinguished by eFluor^450^ labeling and P2X7 expression level were determined by co-staining with an anti-P2X7 antibody (clone RH23A44). Comparative P2X7 451L/P HEK cell analyses were adjusted for P2X7 expression levels by creating a gating region with a comparable P2X7 mean fluorescence intensity (MFI). For the analyses, mixed HEK cells were loaded with Fluo4 and measured in a real-time flow cytometry assay with 2min baseline recording followed by 3 min ATP stimulation (100μM – 2mM). Calcium influx was measured by increase in Fluo4 MFI. **(B)** Analysis of the 500μM ATP P2X7k sample was repeated with a skewed adjustment of P2X7 expression.

**Supplementary figure S3: ARTC2.2 expression is comparable on P2X4ko and WT T cells.** Flow cytometric analyses of cell surface ARTC2.2 expression on CTL, Th and Treg of WT and P2X4ko mice on the B6 and Balb/c background. The mean fluorescence intensity (MFI) of ARTC2 on the different T cell populations from WT and P2X4ko mice (n = 3) was compared.

**Supplementary figure S4: Innate immune cells cultured from bone marrow show reduced P2X7 expression.** P2X7 expression was analyzed on bone marrow-derived dendritic cells (BMDC), macrophages (BMM) and mast cells (BMMC) of B6 and Balb/c WT and P2X4ko mice.

**Supplementary figure S5: mRNA expression of *P2rx4* neighboring genes in innate immune cells of P2X4ko mice.** The expression of the *P2rx4* neighboring genes *Camkk2, ATP2a2, Kdm2b, Anapc5* and *Anapc7* was analyzed in bone marrow-derived dendritic cells (BMDC) and macrophages (BMM) of B6 and Balb/c WT and P2X4ko mice (n = 3-5). Statistical comparison of two groups was performed by using the student’s t test (p < 0.05 = * / p < 0.01 = ** / p < 0.001 = ***, ns = no significant).

