## Supplementary material for "A *P2rx7* passenger mutation affects the vitality and function of immune cells in P2X4ko and other transgenic mice": Table1

| **Gene name** | **Gene start (bp)** | **Gene end (bp)** | **distance to P2rx7 (bp)** | **Gene description** | **129-based KO mice** |
| --- | --- | --- | --- | --- | --- |
| [Rasal1](http://www.ensembl.org/mus_musculus/Gene/Summary?db=core;g=ENSMUSG00000029602) | [120648812](http://www.ensembl.org/mus_musculus/contigview?chr=5&vc_start=120648812&vc_end=120679597) | [120679597](http://www.ensembl.org/mus_musculus/contigview?chr=5&vc_start=120648812&vc_end=120679597) | -1964314 | RAS protein activator like 1 (GAP1 like) [Source:MGI Symbol;Acc:MGI:1330842] | n.a. |
| [Dtx1](http://www.ensembl.org/mus_musculus/Gene/Summary?db=core;g=ENSMUSG00000029603) | [120680202](http://www.ensembl.org/mus_musculus/contigview?chr=5&vc_start=120680202&vc_end=120711927) | [120711927](http://www.ensembl.org/mus_musculus/contigview?chr=5&vc_start=120680202&vc_end=120711927) | -1931984 | deltex 1, E3 ubiquitin ligase [Source:MGI Symbol;Acc:MGI:1352744] | [Dtx1tm1.1Mzl; Dtx1tm1Crey; Dtx1tm1Mjb](http://www.informatics.jax.org/allele/MGI:4355028) |
| [Oas2](http://www.ensembl.org/mus_musculus/Gene/Summary?db=core;g=ENSMUSG00000032690) | [120730333](http://www.ensembl.org/mus_musculus/contigview?chr=5&vc_start=120730333&vc_end=120749853) | [120749853](http://www.ensembl.org/mus_musculus/contigview?chr=5&vc_start=120730333&vc_end=120749853) | -1894058 | 2'-5' oligoadenylate synthetase 2 [Source:MGI Symbol;Acc:MGI:2180852] | [Oas2Gt(OST112989)Lex ;](http://www.informatics.jax.org/allele/MGI:4188821) |
| [Oas3](http://www.ensembl.org/mus_musculus/Gene/Summary?db=core;g=ENSMUSG00000032661) | [120753098](http://www.ensembl.org/mus_musculus/contigview?chr=5&vc_start=120753098&vc_end=120777661) | [120777661](http://www.ensembl.org/mus_musculus/contigview?chr=5&vc_start=120753098&vc_end=120777661) | -1866250 | 2'-5' oligoadenylate synthetase 3 [Source:MGI Symbol;Acc:MGI:2180850] | n.a. |
| [Oas1e](http://www.ensembl.org/mus_musculus/Gene/Summary?db=core;g=ENSMUSG00000066867) | [120786226](http://www.ensembl.org/mus_musculus/contigview?chr=5&vc_start=120786226&vc_end=120795530) | [120795530](http://www.ensembl.org/mus_musculus/contigview?chr=5&vc_start=120786226&vc_end=120795530) | -1848381 | 2'-5' oligoadenylate synthetase 1E [Source:MGI Symbol;Acc:MGI:2180856] | n.a. |
| [Oas1c](http://www.ensembl.org/mus_musculus/Gene/Summary?db=core;g=ENSMUSG00000001166) | [120800194](http://www.ensembl.org/mus_musculus/contigview?chr=5&vc_start=120800194&vc_end=120812514) | [120812514](http://www.ensembl.org/mus_musculus/contigview?chr=5&vc_start=120800194&vc_end=120812514) | -1831397 | 2'-5' oligoadenylate synthetase 1C [Source:MGI Symbol;Acc:MGI:2149633] | n.a. |
| [Oas1b](http://www.ensembl.org/mus_musculus/Gene/Summary?db=core;g=ENSMUSG00000029605) | [120812635](http://www.ensembl.org/mus_musculus/contigview?chr=5&vc_start=120812635&vc_end=120824163) | [120824163](http://www.ensembl.org/mus_musculus/contigview?chr=5&vc_start=120812635&vc_end=120824163) | -1819748 | 2'-5' oligoadenylate synthetase 1B [Source:MGI Symbol;Acc:MGI:97430] | [Oas1btm1.1Brin](http://www.informatics.jax.org/allele/MGI:5285411) |
| [Oas1f](http://www.ensembl.org/mus_musculus/Gene/Summary?db=core;g=ENSMUSG00000053765) | [120847367](http://www.ensembl.org/mus_musculus/contigview?chr=5&vc_start=120847367&vc_end=120857986) | [120857986](http://www.ensembl.org/mus_musculus/contigview?chr=5&vc_start=120847367&vc_end=120857986) | -1785925 | 2'-5' oligoadenylate synthetase 1F [Source:MGI Symbol;Acc:MGI:2180855] | [Oas1fGt(OST425760)Lex](http://www.informatics.jax.org/allele/MGI:4312997) |
| [Oas1h](http://www.ensembl.org/mus_musculus/Gene/Summary?db=core;g=ENSMUSG00000001168) | [120861421](http://www.ensembl.org/mus_musculus/contigview?chr=5&vc_start=120861421&vc_end=120873506) | [120873506](http://www.ensembl.org/mus_musculus/contigview?chr=5&vc_start=120861421&vc_end=120873506) | -1770405 | 2'-5' oligoadenylate synthetase 1H [Source:MGI Symbol;Acc:MGI:2180853] | n.a. |
| [Oas1g](http://www.ensembl.org/mus_musculus/Gene/Summary?db=core;g=ENSMUSG00000066861) | [120876142](http://www.ensembl.org/mus_musculus/contigview?chr=5&vc_start=120876142&vc_end=120887613) | [120887613](http://www.ensembl.org/mus_musculus/contigview?chr=5&vc_start=120876142&vc_end=120887613) | -1756298 | 2'-5' oligoadenylate synthetase 1G [Source:MGI Symbol;Acc:MGI:97429] | n.a. |
| [Oas1a](http://www.ensembl.org/mus_musculus/Gene/Summary?db=core;g=ENSMUSG00000052776) | [120896256](http://www.ensembl.org/mus_musculus/contigview?chr=5&vc_start=120896256&vc_end=120907521) | [120907521](http://www.ensembl.org/mus_musculus/contigview?chr=5&vc_start=120896256&vc_end=120907521) | -1736390 | 2'-5' oligoadenylate synthetase 1A [Source:MGI Symbol;Acc:MGI:2180860] | n.a. |
| [Oas1d](http://www.ensembl.org/mus_musculus/Gene/Summary?db=core;g=ENSMUSG00000032623) | [120914536](http://www.ensembl.org/mus_musculus/contigview?chr=5&vc_start=120914536&vc_end=120921652) | [120921652](http://www.ensembl.org/mus_musculus/contigview?chr=5&vc_start=120914536&vc_end=120921652) | -1722259 | 2'-5' oligoadenylate synthetase 1D [Source:MGI Symbol;Acc:MGI:2140770] | [Oas1dtm1Zuk](http://www.informatics.jax.org/allele/MGI:3581976) |
| [Rph3a](http://www.ensembl.org/mus_musculus/Gene/Summary?db=core;g=ENSMUSG00000029608) | [120940499](http://www.ensembl.org/mus_musculus/contigview?chr=5&vc_start=120940499&vc_end=121010092) | [121010092](http://www.ensembl.org/mus_musculus/contigview?chr=5&vc_start=120940499&vc_end=121010092) | -1633819 | rabphilin 3A [Source:MGI Symbol;Acc:MGI:102788] | [Rph3atm1Sud](http://www.informatics.jax.org/allele/MGI:2179730) |
| [Ptpn11](http://www.ensembl.org/mus_musculus/Gene/Summary?db=core;g=ENSMUSG00000043733) | [121130533](http://www.ensembl.org/mus_musculus/contigview?chr=5&vc_start=121130533&vc_end=121191397) | [121191397](http://www.ensembl.org/mus_musculus/contigview?chr=5&vc_start=121130533&vc_end=121191397) | -1452514 | protein tyrosine phosphatase, non-receptor type 11 [Source:MGI Symbol;Acc:MGI:99511] | [Ptpn11tm1.1Rbns; Ptpn11tm1.1Wbm; Ptpn11tm1Bgn; Ptpn11tm1Rbn; Ptpn11tm1Yan;](http://www.informatics.jax.org/allele/MGI:3852440) |
| [Rpl6](http://www.ensembl.org/mus_musculus/Gene/Summary?db=core;g=ENSMUSG00000029614) | [121204481](http://www.ensembl.org/mus_musculus/contigview?chr=5&vc_start=121204481&vc_end=121209241) | [121209241](http://www.ensembl.org/mus_musculus/contigview?chr=5&vc_start=121204481&vc_end=121209241) | -1434670 | ribosomal protein L6 [Source:MGI Symbol;Acc:MGI:108057] | [Rpl6Gt(OST1622)Lex; Rpl6Gt(PST17838)Mfgc](http://www.informatics.jax.org/allele/MGI:4132022) |
| [Hectd4](http://www.ensembl.org/mus_musculus/Gene/Summary?db=core;g=ENSMUSG00000042744) | [121220219](http://www.ensembl.org/mus_musculus/contigview?chr=5&vc_start=121220219&vc_end=121368577) | [121368577](http://www.ensembl.org/mus_musculus/contigview?chr=5&vc_start=121220219&vc_end=121368577) | -1275334 | HECT domain E3 ubiquitin protein ligase 4 [Source:MGI Symbol;Acc:MGI:3647820] | [Hectd4Gt(255G8)Cmhd; Hectd4Gt(BC0299)Wtsi; Hectd4Gt(BGA536)Byg](http://www.informatics.jax.org/allele/MGI:4960568) |
| [Trafd1](http://www.ensembl.org/mus_musculus/Gene/Summary?db=core;g=ENSMUSG00000042726) | [121371725](http://www.ensembl.org/mus_musculus/contigview?chr=5&vc_start=121371725&vc_end=121385632) | [121385632](http://www.ensembl.org/mus_musculus/contigview?chr=5&vc_start=121371725&vc_end=121385632) | -1258279 | TRAF type zinc finger domain containing 1 [Source:MGI Symbol;Acc:MGI:1923551] | [Trafd1tm1Ayos](http://www.informatics.jax.org/allele/MGI:3831676) |
| [Naa25](http://www.ensembl.org/mus_musculus/Gene/Summary?db=core;g=ENSMUSG00000042719) | [121397936](http://www.ensembl.org/mus_musculus/contigview?chr=5&vc_start=121397936&vc_end=121444378) | [121444378](http://www.ensembl.org/mus_musculus/contigview?chr=5&vc_start=121397936&vc_end=121444378) | -1199533 | N(alpha)-acetyltransferase 25, NatB auxiliary subunit [Source:MGI Symbol;Acc:MGI:2442563] | [Naa25Gt(RRK280)Byg; Naa25Gt(AL0004)Wtsi](http://www.informatics.jax.org/allele/MGI:4130258) |
| [Erp29](http://www.ensembl.org/mus_musculus/Gene/Summary?db=core;g=ENSMUSG00000029616) | [121428590](http://www.ensembl.org/mus_musculus/contigview?chr=5&vc_start=121428590&vc_end=121452506) | [121452506](http://www.ensembl.org/mus_musculus/contigview?chr=5&vc_start=121428590&vc_end=121452506) | -1191405 | endoplasmic reticulum protein 29 [Source:MGI Symbol;Acc:MGI:1914647] | [Erp29tm1Dfer; Erp29Gt(KST171)Byg; Erp29Gt(G014A01)Wrst](http://www.informatics.jax.org/allele/MGI:6342880) |
| [Tmem116](http://www.ensembl.org/mus_musculus/Gene/Summary?db=core;g=ENSMUSG00000029452) | [121451893](http://www.ensembl.org/mus_musculus/contigview?chr=5&vc_start=121451893&vc_end=121524183) | [121524183](http://www.ensembl.org/mus_musculus/contigview?chr=5&vc_start=121451893&vc_end=121524183) | -1119728 | transmembrane protein 116 [Source:MGI Symbol;Acc:MGI:1924712] | [Tmem116Gt(OST44984)Lex; Tmem116Gt(PST12765)Mfgc](http://www.informatics.jax.org/allele/MGI:4159310) |
| [Adam1b](http://www.ensembl.org/mus_musculus/Gene/Summary?db=core;g=ENSMUSG00000062438) | [121500098](http://www.ensembl.org/mus_musculus/contigview?chr=5&vc_start=121500098&vc_end=121503435) | [121503435](http://www.ensembl.org/mus_musculus/contigview?chr=5&vc_start=121500098&vc_end=121503435) | -1140476 | a disintegrin and metallopeptidase domain 1b [Source:MGI Symbol;Acc:MGI:2429506] | n.a. |
| [Adam1a](http://www.ensembl.org/mus_musculus/Gene/Summary?db=core;g=ENSMUSG00000072647) | [121518576](http://www.ensembl.org/mus_musculus/contigview?chr=5&vc_start=121518576&vc_end=121545482) | [121545482](http://www.ensembl.org/mus_musculus/contigview?chr=5&vc_start=121518576&vc_end=121545482) | -1098429 | a disintegrin and metallopeptidase domain 1a [Source:MGI Symbol;Acc:MGI:2429504] | [Adam1btm1Tba;](http://www.informatics.jax.org/allele/MGI:3628759) |
| [Mapkapk5](http://www.ensembl.org/mus_musculus/Gene/Summary?db=core;g=ENSMUSG00000029454) | [121525038](http://www.ensembl.org/mus_musculus/contigview?chr=5&vc_start=121525038&vc_end=121545905) | [121545905](http://www.ensembl.org/mus_musculus/contigview?chr=5&vc_start=121525038&vc_end=121545905) | -1098006 | MAP kinase-activated protein kinase 5 [Source:MGI Symbol;Acc:MGI:1333110] | [Mapkapk5tm1Mgl; Mapkapk5tm1Pqs](http://www.informatics.jax.org/allele/MGI:2680287) |
| [Aldh2](http://www.ensembl.org/mus_musculus/Gene/Summary?db=core;g=ENSMUSG00000029455) | [121566027](http://www.ensembl.org/mus_musculus/contigview?chr=5&vc_start=121566027&vc_end=121593824) | [121593824](http://www.ensembl.org/mus_musculus/contigview?chr=5&vc_start=121566027&vc_end=121593824) | -1050087 | aldehyde dehydrogenase 2, mitochondrial [Source:MGI Symbol;Acc:MGI:99600] | [Aldh2Gt(OST7285)Lex; Aldh2Gt(D053A10)Wrst;](http://www.informatics.jax.org/allele/MGI:3530243) |
| [Acad12](http://www.ensembl.org/mus_musculus/Gene/Summary?db=core;g=ENSMUSG00000042647) | [121596775](http://www.ensembl.org/mus_musculus/contigview?chr=5&vc_start=121596775&vc_end=121618938) | [121618938](http://www.ensembl.org/mus_musculus/contigview?chr=5&vc_start=121596775&vc_end=121618938) | -1024973 | acyl-Coenzyme A dehydrogenase family, member 12 [Source:MGI Symbol;Acc:MGI:2443320] | [Acad12Gt(OST300106)Lex](http://www.informatics.jax.org/allele/MGI:4270848) |
| [Acad10](http://www.ensembl.org/mus_musculus/Gene/Summary?db=core;g=ENSMUSG00000029456) | [121621026](http://www.ensembl.org/mus_musculus/contigview?chr=5&vc_start=121621026&vc_end=121660514) | [121660514](http://www.ensembl.org/mus_musculus/contigview?chr=5&vc_start=121621026&vc_end=121660514) | -983397 | acyl-Coenzyme A dehydrogenase family, member 10 [Source:MGI Symbol;Acc:MGI:1919235] | [Acad10Gt(OST448289)Lex](http://www.informatics.jax.org/allele/MGI:4320804) |
| [Brap](http://www.ensembl.org/mus_musculus/Gene/Summary?db=core;g=ENSMUSG00000029458) | [121660563](http://www.ensembl.org/mus_musculus/contigview?chr=5&vc_start=121660563&vc_end=121687256) | [121687256](http://www.ensembl.org/mus_musculus/contigview?chr=5&vc_start=121660563&vc_end=121687256) | -956655 | BRCA1 associated protein [Source:MGI Symbol;Acc:MGI:1919649] | n.a. |
| [Atxn2](http://www.ensembl.org/mus_musculus/Gene/Summary?db=core;g=ENSMUSG00000042605) | [121711337](http://www.ensembl.org/mus_musculus/contigview?chr=5&vc_start=121711337&vc_end=121816493) | [121816493](http://www.ensembl.org/mus_musculus/contigview?chr=5&vc_start=121711337&vc_end=121816493) | -827418 | ataxin 2 [Source:MGI Symbol;Acc:MGI:1277223] | [Atxn2tm1.1Geno ; Atxn2tm1Plt ; Atxn2tm2.1Geno](http://www.informatics.jax.org/allele/summary?markerId=MGI:1277223&alleleType=Targeted) |
| [Sh2b3](http://www.ensembl.org/mus_musculus/Gene/Summary?db=core;g=ENSMUSG00000042594) | [121815488](http://www.ensembl.org/mus_musculus/contigview?chr=5&vc_start=121815488&vc_end=121837646) | [121837646](http://www.ensembl.org/mus_musculus/contigview?chr=5&vc_start=121815488&vc_end=121837646) | -806265 | SH2B adaptor protein 3 [Source:MGI Symbol;Acc:MGI:893598] | [Sh2b3tm1Paw ; Sh2b3tm1Rmp](http://www.informatics.jax.org/allele/summary?markerId=MGI:893598&alleleType=Targeted) |
| [Pheta1](http://www.ensembl.org/mus_musculus/Gene/Summary?db=core;g=ENSMUSG00000044134) | [121848984](http://www.ensembl.org/mus_musculus/contigview?chr=5&vc_start=121848984&vc_end=121854632) | [121854632](http://www.ensembl.org/mus_musculus/contigview?chr=5&vc_start=121848984&vc_end=121854632) | -789279 | PH domain containing endocytic trafficking adaptor 1 [Source:MGI Symbol;Acc:MGI:2442708] | n.a |
| [Cux2](http://www.ensembl.org/mus_musculus/Gene/Summary?db=core;g=ENSMUSG00000042589) | [121856366](http://www.ensembl.org/mus_musculus/contigview?chr=5&vc_start=121856366&vc_end=122050102) | [122050102](http://www.ensembl.org/mus_musculus/contigview?chr=5&vc_start=121856366&vc_end=122050102) | -593809 | cut-like homeobox 2 [Source:MGI Symbol;Acc:MGI:107321] | [Cux2tm1.1Nieto](http://www.informatics.jax.org/allele/MGI:4429541) |
| [Myl2](http://www.ensembl.org/mus_musculus/Gene/Summary?db=core;g=ENSMUSG00000013936) | [122100951](http://www.ensembl.org/mus_musculus/contigview?chr=5&vc_start=122100951&vc_end=122113472) | [122113472](http://www.ensembl.org/mus_musculus/contigview?chr=5&vc_start=122100951&vc_end=122113472) | -530439 | myosin, light polypeptide 2, regulatory, cardiac, slow [Source:MGI Symbol;Acc:MGI:97272] | [Myl2tm1(cre)Krc ; Myl2tm1(Hand1)Tana ; Myl2tm1.1Chen ; Myl2tm2.1Chen](http://www.informatics.jax.org/allele/summary?markerId=MGI:97272&alleleType=Targeted) |
| [Ccdc63](http://www.ensembl.org/mus_musculus/Gene/Summary?db=core;g=ENSMUSG00000043036) | [122108040](http://www.ensembl.org/mus_musculus/contigview?chr=5&vc_start=122108040&vc_end=122140823) | [122140823](http://www.ensembl.org/mus_musculus/contigview?chr=5&vc_start=122108040&vc_end=122140823) | -503088 | coiled-coil domain containing 63 [Source:MGI Symbol;Acc:MGI:3607777] | n.a |
| [Ppp1cc](http://www.ensembl.org/mus_musculus/Gene/Summary?db=core;g=ENSMUSG00000004455) | [122158278](http://www.ensembl.org/mus_musculus/contigview?chr=5&vc_start=122158278&vc_end=122175273) | [122175273](http://www.ensembl.org/mus_musculus/contigview?chr=5&vc_start=122158278&vc_end=122175273) | -468638 | protein phosphatase 1 catalytic subunit gamma [Source:MGI Symbol;Acc:MGI:104872] | [Ppp1cctm1Lex ; Ppp1cctm1Var](http://www.informatics.jax.org/allele/summary?markerId=MGI:104872&alleleType=Targeted) |
| [Hvcn1](http://www.ensembl.org/mus_musculus/Gene/Summary?db=core;g=ENSMUSG00000064267) | [122206804](http://www.ensembl.org/mus_musculus/contigview?chr=5&vc_start=122206804&vc_end=122242297) | [122242297](http://www.ensembl.org/mus_musculus/contigview?chr=5&vc_start=122206804&vc_end=122242297) | -401614 | hydrogen voltage-gated channel 1 [Source:MGI Symbol;Acc:MGI:1921346] | [Hvcn1Gt(RRN293)Byg](http://www.informatics.jax.org/allele/MGI:3843777) |
| [Tctn1](http://www.ensembl.org/mus_musculus/Gene/Summary?db=core;g=ENSMUSG00000038593) | [122237848](http://www.ensembl.org/mus_musculus/contigview?chr=5&vc_start=122237848&vc_end=122264460) | [122264460](http://www.ensembl.org/mus_musculus/contigview?chr=5&vc_start=122237848&vc_end=122264460) | -379451 | tectonic family member 1 [Source:MGI Symbol;Acc:MGI:3603820] | n.a |
| [Pptc7](http://www.ensembl.org/mus_musculus/Gene/Summary?db=core;g=ENSMUSG00000038582) | [122284365](http://www.ensembl.org/mus_musculus/contigview?chr=5&vc_start=122284365&vc_end=122324281) | [122324281](http://www.ensembl.org/mus_musculus/contigview?chr=5&vc_start=122284365&vc_end=122324281) | -319630 | PTC7 protein phosphatase homolog [Source:MGI Symbol;Acc:MGI:2444593] | n.a |
| [Rad9b](http://www.ensembl.org/mus_musculus/Gene/Summary?db=core;g=ENSMUSG00000038569) | [122323223](http://www.ensembl.org/mus_musculus/contigview?chr=5&vc_start=122323223&vc_end=122354233) | [122354233](http://www.ensembl.org/mus_musculus/contigview?chr=5&vc_start=122323223&vc_end=122354233) | -289678 | RAD9 checkpoint clamp component B [Source:MGI Symbol;Acc:MGI:2385231] | [Rad9btm1Lieb](http://www.informatics.jax.org/allele/MGI:3526236) |
| [Vps29](http://www.ensembl.org/mus_musculus/Gene/Summary?db=core;g=ENSMUSG00000029462) | [122354369](http://www.ensembl.org/mus_musculus/contigview?chr=5&vc_start=122354369&vc_end=122364984) | [122364984](http://www.ensembl.org/mus_musculus/contigview?chr=5&vc_start=122354369&vc_end=122364984) | -278927 | VPS29 retromer complex component [Source:MGI Symbol;Acc:MGI:1928344] | [Vps29Gt(OST309649)Lex ; Vps29Gt(OST17630)Lex ; Vps29Gt(OST18210)Lex ; Vps29Gt(OST24044)Lex ; Vps29Gt(OST28280)Lex ; Vps29Gt(OST93118)Lex ; Vps29Gt(OST98823)Lex ; Vps29Gt(OST113533)Lex ; Vps29Gt(OST127837)Lex ; Vps29Gt(OST135740)Lex ; Vps29Gt(OST456377)Lex](http://www.informatics.jax.org/allele/summary?markerId=MGI:1928344&alleleType=Gene%20trapped) |
| [Fam216a](http://www.ensembl.org/mus_musculus/Gene/Summary?db=core;g=ENSMUSG00000029463) | [122364580](http://www.ensembl.org/mus_musculus/contigview?chr=5&vc_start=122364580&vc_end=122372364) | [122372364](http://www.ensembl.org/mus_musculus/contigview?chr=5&vc_start=122364580&vc_end=122372364) | -271547 | family with sequence similarity 216, member A [Source:MGI Symbol;Acc:MGI:1916198] | n.a |
| [Gpn3](http://www.ensembl.org/mus_musculus/Gene/Summary?db=core;g=ENSMUSG00000029464) | [122371876](http://www.ensembl.org/mus_musculus/contigview?chr=5&vc_start=122371876&vc_end=122382902) | [122382902](http://www.ensembl.org/mus_musculus/contigview?chr=5&vc_start=122371876&vc_end=122382902) | -261009 | GPN-loop GTPase 3 [Source:MGI Symbol;Acc:MGI:1289326] | n.a |
| [Arpc3](http://www.ensembl.org/mus_musculus/Gene/Summary?db=core;g=ENSMUSG00000029465) | [122391878](http://www.ensembl.org/mus_musculus/contigview?chr=5&vc_start=122391878&vc_end=122414184) | [122414184](http://www.ensembl.org/mus_musculus/contigview?chr=5&vc_start=122391878&vc_end=122414184) | -229727 | actin related protein 2/3 complex, subunit 3 [Source:MGI Symbol;Acc:MGI:1928375] | [Arpc3tm1Jtak](http://www.informatics.jax.org/allele/MGI:3664649) |
| [Anapc7](http://www.ensembl.org/mus_musculus/Gene/Summary?db=core;g=ENSMUSG00000029466) | [122421693](http://www.ensembl.org/mus_musculus/contigview?chr=5&vc_start=122421693&vc_end=122444912) | [122444912](http://www.ensembl.org/mus_musculus/contigview?chr=5&vc_start=122421693&vc_end=122444912) | -198999 | anaphase promoting complex subunit 7 [Source:MGI Symbol;Acc:MGI:1929711] | n.a |
| [Atp2a2](http://www.ensembl.org/mus_musculus/Gene/Summary?db=core;g=ENSMUSG00000029467) | [122453513](http://www.ensembl.org/mus_musculus/contigview?chr=5&vc_start=122453513&vc_end=122502225) | [122502225](http://www.ensembl.org/mus_musculus/contigview?chr=5&vc_start=122453513&vc_end=122502225) | -141686 | ATPase, Ca++ transporting, cardiac muscle, slow twitch 2 [Source:MGI Symbol;Acc:MGI:88110] | [Atp2a2tm1.1Iemr ; Atp2a2tm1Fwuy ; Atp2a2tm1Ges](http://www.informatics.jax.org/allele/summary?markerId=MGI:88110&alleleType=Targeted) |
| [Ift81](http://www.ensembl.org/mus_musculus/Gene/Summary?db=core;g=ENSMUSG00000029469) | [122550204](http://www.ensembl.org/mus_musculus/contigview?chr=5&vc_start=122550204&vc_end=122614518) | [122614518](http://www.ensembl.org/mus_musculus/contigview?chr=5&vc_start=122550204&vc_end=122614518) | -29393 | intraflagellar transport 81 [Source:MGI Symbol;Acc:MGI:1098597] | n.a |
| [**P2rx7**](http://www.ensembl.org/mus_musculus/Gene/Summary?db=core;g=ENSMUSG00000029468) | [**122643911**](http://www.ensembl.org/mus_musculus/contigview?chr=5&vc_start=122643911&vc_end=122691432) | [**122691432**](http://www.ensembl.org/mus_musculus/contigview?chr=5&vc_start=122643911&vc_end=122691432) | **0** | **purinergic receptor P2X, ligand-gated ion channel, 7 [Source:MGI Symbol;Acc:MGI:1339957]** |  |
| [P2rx4](http://www.ensembl.org/mus_musculus/Gene/Summary?db=core;g=ENSMUSG00000029470) | [122707544](http://www.ensembl.org/mus_musculus/contigview?chr=5&vc_start=122707544&vc_end=122729738) | [122729738](http://www.ensembl.org/mus_musculus/contigview?chr=5&vc_start=122707544&vc_end=122729738) | 16112 | purinergic receptor P2X, ligand-gated ion channel 4 [Source:MGI Symbol;Acc:MGI:1338859] | [P2rx4tm1Rass ; P2rx4tm1Dgen](http://www.informatics.jax.org/allele/MGI:3665297) |
| [Camkk2](http://www.ensembl.org/mus_musculus/Gene/Summary?db=core;g=ENSMUSG00000029471) | [122731170](http://www.ensembl.org/mus_musculus/contigview?chr=5&vc_start=122731170&vc_end=122779409) | [122779409](http://www.ensembl.org/mus_musculus/contigview?chr=5&vc_start=122731170&vc_end=122779409) | 39738 | calcium/calmodulin-dependent protein kinase kinase 2, beta [Source:MGI Symbol;Acc:MGI:2444812] | [Camkk2tm1Kpg ; Camkk2tm1Shyy ; Camkk2tm1Tch ; Camkk2tm2.1Kpg ; Camkk2tm2Kpg](http://www.informatics.jax.org/allele/summary?markerId=MGI:2444812&alleleType=Targeted) |
| [Anapc5](http://www.ensembl.org/mus_musculus/Gene/Summary?db=core;g=ENSMUSG00000029472) | [122787459](http://www.ensembl.org/mus_musculus/contigview?chr=5&vc_start=122787459&vc_end=122821339) | [122821339](http://www.ensembl.org/mus_musculus/contigview?chr=5&vc_start=122787459&vc_end=122821339) | 96027 | anaphase-promoting complex subunit 5 [Source:MGI Symbol;Acc:MGI:1929722] | n.a |
| [Rnf34](http://www.ensembl.org/mus_musculus/Gene/Summary?db=core;g=ENSMUSG00000029474) | [122850048](http://www.ensembl.org/mus_musculus/contigview?chr=5&vc_start=122850048&vc_end=122871291) | [122871291](http://www.ensembl.org/mus_musculus/contigview?chr=5&vc_start=122850048&vc_end=122871291) | 158616 | ring finger protein 34 [Source:MGI Symbol;Acc:MGI:2153340] | n.a |
| [Kdm2b](http://www.ensembl.org/mus_musculus/Gene/Summary?db=core;g=ENSMUSG00000029475) | [122870665](http://www.ensembl.org/mus_musculus/contigview?chr=5&vc_start=122870665&vc_end=122989823) | [122989823](http://www.ensembl.org/mus_musculus/contigview?chr=5&vc_start=122870665&vc_end=122989823) | 179233 | lysine (K)-specific demethylase 2B [Source:MGI Symbol;Acc:MGI:1354737] | [Col1a1tm1(CMV/tetO-Kdm2b)Atz ; Kdm2btm1.1Atz ; Kdm2btm1.1Bes ; Kdm2btm1Nobu](http://www.informatics.jax.org/allele/summary?markerId=MGI:1354737&alleleType=Targeted) |
| [Orai1](http://www.ensembl.org/mus_musculus/Gene/Summary?db=core;g=ENSMUSG00000049686) | [123015074](http://www.ensembl.org/mus_musculus/contigview?chr=5&vc_start=123015074&vc_end=123030456) | [123030456](http://www.ensembl.org/mus_musculus/contigview?chr=5&vc_start=123015074&vc_end=123030456) | 323642 | ORAI calcium release-activated calcium modulator 1 [Source:MGI Symbol;Acc:MGI:1925542] | [Orai1tm1.1Ygwa](http://www.informatics.jax.org/allele/MGI:6149978) |
| [Morn3](http://www.ensembl.org/mus_musculus/Gene/Summary?db=core;g=ENSMUSG00000029477) | [123035769](http://www.ensembl.org/mus_musculus/contigview?chr=5&vc_start=123035769&vc_end=123047016) | [123047016](http://www.ensembl.org/mus_musculus/contigview?chr=5&vc_start=123035769&vc_end=123047016) | 344337 | MORN repeat containing 3 [Source:MGI Symbol;Acc:MGI:1922140] | n.a |
| [Tmem120b](http://www.ensembl.org/mus_musculus/Gene/Summary?db=core;g=ENSMUSG00000054434) | [123068415](http://www.ensembl.org/mus_musculus/contigview?chr=5&vc_start=123068415&vc_end=123117749) | [123117749](http://www.ensembl.org/mus_musculus/contigview?chr=5&vc_start=123068415&vc_end=123117749) | 376983 | transmembrane protein 120B [Source:MGI Symbol;Acc:MGI:3603158] | n.a |
| [Rhof](http://www.ensembl.org/mus_musculus/Gene/Summary?db=core;g=ENSMUSG00000029449) | [123103044](http://www.ensembl.org/mus_musculus/contigview?chr=5&vc_start=123103044&vc_end=123132692) | [123132692](http://www.ensembl.org/mus_musculus/contigview?chr=5&vc_start=123103044&vc_end=123132692) | 411612 | ras homolog family member F (in filopodia) [Source:MGI Symbol;Acc:MGI:1345629] | n.a |
| [Setd1b](http://www.ensembl.org/mus_musculus/Gene/Summary?db=core;g=ENSMUSG00000038384) | [123142193](http://www.ensembl.org/mus_musculus/contigview?chr=5&vc_start=123142193&vc_end=123168629) | [123168629](http://www.ensembl.org/mus_musculus/contigview?chr=5&vc_start=123142193&vc_end=123168629) | 450761 | SET domain containing 1B [Source:MGI Symbol;Acc:MGI:2652820] | [Setd1btm1.1Afst ; Setd1btm1Afst](http://www.informatics.jax.org/allele/summary?markerId=MGI:2652820&alleleType=Targeted) |
| [Psmd9](http://www.ensembl.org/mus_musculus/Gene/Summary?db=core;g=ENSMUSG00000029440) | [123169413](http://www.ensembl.org/mus_musculus/contigview?chr=5&vc_start=123169413&vc_end=123250131) | [123250131](http://www.ensembl.org/mus_musculus/contigview?chr=5&vc_start=123169413&vc_end=123250131) | 477981 | proteasome (prosome, macropain) 26S subunit, non-ATPase, 9 [Source:MGI Symbol;Acc:MGI:1914401] | n.a |
| [Hpd](http://www.ensembl.org/mus_musculus/Gene/Summary?db=core;g=ENSMUSG00000029445) | [123171807](http://www.ensembl.org/mus_musculus/contigview?chr=5&vc_start=123171807&vc_end=123182727) | [123182727](http://www.ensembl.org/mus_musculus/contigview?chr=5&vc_start=123171807&vc_end=123182727) | 480375 | 4-hydroxyphenylpyruvic acid dioxygenase [Source:MGI Symbol;Acc:MGI:96213] | n.a |
| [Wdr66](http://www.ensembl.org/mus_musculus/Gene/Summary?db=core;g=ENSMUSG00000029442) | [123252102](http://www.ensembl.org/mus_musculus/contigview?chr=5&vc_start=123252102&vc_end=123327484) | [123327484](http://www.ensembl.org/mus_musculus/contigview?chr=5&vc_start=123252102&vc_end=123327484) | 560670 | WD repeat domain 66 [Source:MGI Symbol;Acc:MGI:1918495] | n.a |
| [Bcl7a](http://www.ensembl.org/mus_musculus/Gene/Summary?db=core;g=ENSMUSG00000029438) | [123343834](http://www.ensembl.org/mus_musculus/contigview?chr=5&vc_start=123343834&vc_end=123374992) | [123374992](http://www.ensembl.org/mus_musculus/contigview?chr=5&vc_start=123343834&vc_end=123374992) | 652402 | B cell CLL/lymphoma 7A [Source:MGI Symbol;Acc:MGI:1924295] | n.a |
| [Mlxip](http://www.ensembl.org/mus_musculus/Gene/Summary?db=core;g=ENSMUSG00000038342) | [123394798](http://www.ensembl.org/mus_musculus/contigview?chr=5&vc_start=123394798&vc_end=123457932) | [123457932](http://www.ensembl.org/mus_musculus/contigview?chr=5&vc_start=123394798&vc_end=123457932) | 703366 | MLX interacting protein [Source:MGI Symbol;Acc:MGI:2141183] | [Mlxiptm1.1Lchan ; Mlxiptm1.2Lchan](http://www.informatics.jax.org/allele/summary?markerId=MGI:2141183&alleleType=Targeted) |
| [Rpl31-ps6](http://www.ensembl.org/mus_musculus/Gene/Summary?db=core;g=ENSMUSG00000082532) | [123466509](http://www.ensembl.org/mus_musculus/contigview?chr=5&vc_start=123466509&vc_end=123466886) | [123466886](http://www.ensembl.org/mus_musculus/contigview?chr=5&vc_start=123466509&vc_end=123466886) | 775077 | ribosomal protein L31, pseudogene 6 [Source:MGI Symbol;Acc:MGI:3783190] | n.a |
| [Il31](http://www.ensembl.org/mus_musculus/Gene/Summary?db=core;g=ENSMUSG00000029437) | [123480157](http://www.ensembl.org/mus_musculus/contigview?chr=5&vc_start=123480157&vc_end=123489489) | [123489489](http://www.ensembl.org/mus_musculus/contigview?chr=5&vc_start=123480157&vc_end=123489489) | 788725 | interleukin 31 [Source:MGI Symbol;Acc:MGI:1923649] | n.a |
| [Lrrc43](http://www.ensembl.org/mus_musculus/Gene/Summary?db=core;g=ENSMUSG00000063409) | [123489305](http://www.ensembl.org/mus_musculus/contigview?chr=5&vc_start=123489305&vc_end=123508205) | [123508205](http://www.ensembl.org/mus_musculus/contigview?chr=5&vc_start=123489305&vc_end=123508205) | 797873 | leucine rich repeat containing 43 [Source:MGI Symbol;Acc:MGI:2685907] | n.a |
| [Diablo](http://www.ensembl.org/mus_musculus/Gene/Summary?db=core;g=ENSMUSG00000029433) | [123509765](http://www.ensembl.org/mus_musculus/contigview?chr=5&vc_start=123509765&vc_end=123524176) | [123524176](http://www.ensembl.org/mus_musculus/contigview?chr=5&vc_start=123509765&vc_end=123524176) | 818333 | diablo, IAP-binding mitochondrial protein [Source:MGI Symbol;Acc:MGI:1913843] | [Diablotm1Mak](http://www.informatics.jax.org/allele/MGI:2183277) |
| [B3gnt4](http://www.ensembl.org/mus_musculus/Gene/Summary?db=core;g=ENSMUSG00000029431) | [123510460](http://www.ensembl.org/mus_musculus/contigview?chr=5&vc_start=123510460&vc_end=123511882) | [123511882](http://www.ensembl.org/mus_musculus/contigview?chr=5&vc_start=123510460&vc_end=123511882) | 819028 | UDP-GlcNAc:betaGal beta-1,3-N-acetylglucosaminyltransferase 4 [Source:MGI Symbol;Acc:MGI:2680208] | n.a |
| [Vps33a](http://www.ensembl.org/mus_musculus/Gene/Summary?db=core;g=ENSMUSG00000029434) | [123528659](http://www.ensembl.org/mus_musculus/contigview?chr=5&vc_start=123528659&vc_end=123573038) | [123573038](http://www.ensembl.org/mus_musculus/contigview?chr=5&vc_start=123528659&vc_end=123573038) | 837227 | VPS33A CORVET/HOPS core subunit [Source:MGI Symbol;Acc:MGI:1924823] | n.a |
| [Clip1](http://www.ensembl.org/mus_musculus/Gene/Summary?db=core;g=ENSMUSG00000049550) | [123577795](http://www.ensembl.org/mus_musculus/contigview?chr=5&vc_start=123577795&vc_end=123684618) | [123684618](http://www.ensembl.org/mus_musculus/contigview?chr=5&vc_start=123577795&vc_end=123684618) | 886363 | CAP-GLY domain containing linker protein 1 [Source:MGI Symbol;Acc:MGI:1928401] | [Clip1tm1.1Gal ; Clip1tm1Gal](http://www.informatics.jax.org/allele/summary?markerId=MGI:1928401&alleleType=Targeted) |
| [Zcchc8](http://www.ensembl.org/mus_musculus/Gene/Summary?db=core;g=ENSMUSG00000029427) | [123698294](http://www.ensembl.org/mus_musculus/contigview?chr=5&vc_start=123698294&vc_end=123721100) | [123721100](http://www.ensembl.org/mus_musculus/contigview?chr=5&vc_start=123698294&vc_end=123721100) | 1006862 | zinc finger, CCHC domain containing 8 [Source:MGI Symbol;Acc:MGI:1917900] | n.a |
| [Rsrc2](http://www.ensembl.org/mus_musculus/Gene/Summary?db=core;g=ENSMUSG00000029422) | [123728426](http://www.ensembl.org/mus_musculus/contigview?chr=5&vc_start=123728426&vc_end=123749414) | [123749414](http://www.ensembl.org/mus_musculus/contigview?chr=5&vc_start=123728426&vc_end=123749414) | 1036994 | arginine/serine-rich coiled-coil 2 [Source:MGI Symbol;Acc:MGI:1913489] | n.a |
| [Kntc1](http://www.ensembl.org/mus_musculus/Gene/Summary?db=core;g=ENSMUSG00000029414) | [123749716](http://www.ensembl.org/mus_musculus/contigview?chr=5&vc_start=123749716&vc_end=123821593) | [123821593](http://www.ensembl.org/mus_musculus/contigview?chr=5&vc_start=123749716&vc_end=123821593) | 1058284 | kinetochore associated 1 [Source:MGI Symbol;Acc:MGI:2673709] | [Kntc1Gt(OST40060)Lex](http://www.informatics.jax.org/allele/MGI:4156696) |
| [Hcar2](http://www.ensembl.org/mus_musculus/Gene/Summary?db=core;g=ENSMUSG00000045502) | [123863570](http://www.ensembl.org/mus_musculus/contigview?chr=5&vc_start=123863570&vc_end=123865499) | [123865499](http://www.ensembl.org/mus_musculus/contigview?chr=5&vc_start=123863570&vc_end=123865499) | 1172138 | hydroxycarboxylic acid receptor 2 [Source:MGI Symbol;Acc:MGI:1933383] | [Hcar2tm1Lex; Hcar2tm1Soff](http://www.informatics.jax.org/allele/MGI:3528952) |
| [Hcar1](http://www.ensembl.org/mus_musculus/Gene/Summary?db=core;g=ENSMUSG00000049241) | [123876736](http://www.ensembl.org/mus_musculus/contigview?chr=5&vc_start=123876736&vc_end=123880020) | [123880020](http://www.ensembl.org/mus_musculus/contigview?chr=5&vc_start=123876736&vc_end=123880020) | 1185304 | hydrocarboxylic acid receptor 1 [Source:MGI Symbol;Acc:MGI:2441671] | n.a |
| [Denr](http://www.ensembl.org/mus_musculus/Gene/Summary?db=core;g=ENSMUSG00000023106) | [123907175](http://www.ensembl.org/mus_musculus/contigview?chr=5&vc_start=123907175&vc_end=123928835) | [123928835](http://www.ensembl.org/mus_musculus/contigview?chr=5&vc_start=123907175&vc_end=123928835) | 1215743 | density-regulated protein [Source:MGI Symbol;Acc:MGI:1915434] | [DenrGt(CSI770)Byg; DenrGt(OST114)Lex](http://www.informatics.jax.org/allele/MGI:3880269) |
| [Ccdc62](http://www.ensembl.org/mus_musculus/Gene/Summary?db=core;g=ENSMUSG00000061882) | [123930679](http://www.ensembl.org/mus_musculus/contigview?chr=5&vc_start=123930679&vc_end=123969895) | [123969895](http://www.ensembl.org/mus_musculus/contigview?chr=5&vc_start=123930679&vc_end=123969895) | 1239247 | coiled-coil domain containing 62 [Source:MGI Symbol;Acc:MGI:2684996] | n.a |
| [Hip1r](http://www.ensembl.org/mus_musculus/Gene/Summary?db=core;g=ENSMUSG00000000915) | [123973628](http://www.ensembl.org/mus_musculus/contigview?chr=5&vc_start=123973628&vc_end=124005558) | [124005558](http://www.ensembl.org/mus_musculus/contigview?chr=5&vc_start=123973628&vc_end=124005558) | 1282196 | huntingtin interacting protein 1 related [Source:MGI Symbol;Acc:MGI:1352504] | [Hip1rtm1Tsr](http://www.informatics.jax.org/allele/MGI:3045594) |
| [Vps37b](http://www.ensembl.org/mus_musculus/Gene/Summary?db=core;g=ENSMUSG00000066278) | [124004641](http://www.ensembl.org/mus_musculus/contigview?chr=5&vc_start=124004641&vc_end=124032270) | [124032270](http://www.ensembl.org/mus_musculus/contigview?chr=5&vc_start=124004641&vc_end=124032270) | 1313209 | vacuolar protein sorting 37B [Source:MGI Symbol;Acc:MGI:1916724] | n.a |
| [Abcb9](http://www.ensembl.org/mus_musculus/Gene/Summary?db=core;g=ENSMUSG00000029408) | [124061530](http://www.ensembl.org/mus_musculus/contigview?chr=5&vc_start=124061530&vc_end=124095798) | [124095798](http://www.ensembl.org/mus_musculus/contigview?chr=5&vc_start=124061530&vc_end=124095798) | 1370098 | ATP-binding cassette, sub-family B (MDR/TAP), member 9 [Source:MGI Symbol;Acc:MGI:1861729] | [Abcb9Gt(231G11)Cmhd](http://www.informatics.jax.org/allele/MGI:4969016) |
| [Ogfod2](http://www.ensembl.org/mus_musculus/Gene/Summary?db=core;g=ENSMUSG00000023707) | [124112297](http://www.ensembl.org/mus_musculus/contigview?chr=5&vc_start=124112297&vc_end=124115483) | [124115483](http://www.ensembl.org/mus_musculus/contigview?chr=5&vc_start=124112297&vc_end=124115483) | 1420865 | 2-oxoglutarate and iron-dependent oxygenase domain containing 2 [Source:MGI Symbol;Acc:MGI:1913877] | n.a |
| [Arl6ip4](http://www.ensembl.org/mus_musculus/Gene/Summary?db=core;g=ENSMUSG00000029404) | [124116089](http://www.ensembl.org/mus_musculus/contigview?chr=5&vc_start=124116089&vc_end=124118196) | [124118196](http://www.ensembl.org/mus_musculus/contigview?chr=5&vc_start=124116089&vc_end=124118196) | 1424657 | ADP-ribosylation factor-like 6 interacting protein 4 [Source:MGI Symbol;Acc:MGI:1929500] | [Arl6ip4Gt(OST31151)Lex](http://www.informatics.jax.org/allele/MGI:4151141) |
| [Pitpnm2](http://www.ensembl.org/mus_musculus/Gene/Summary?db=core;g=ENSMUSG00000029406) | [124118690](http://www.ensembl.org/mus_musculus/contigview?chr=5&vc_start=124118690&vc_end=124249760) | [124249760](http://www.ensembl.org/mus_musculus/contigview?chr=5&vc_start=124118690&vc_end=124249760) | 1427258 | phosphatidylinositol transfer protein, membrane-associated 2 [Source:MGI Symbol;Acc:MGI:1336192] | [Pitpnm2tm1Tili](http://www.informatics.jax.org/allele/MGI:2679326) |
| [Pitpnm2os2](http://www.ensembl.org/mus_musculus/Gene/Summary?db=core;g=ENSMUSG00000089842) | [124194907](http://www.ensembl.org/mus_musculus/contigview?chr=5&vc_start=124194907&vc_end=124200346) | [124200346](http://www.ensembl.org/mus_musculus/contigview?chr=5&vc_start=124194907&vc_end=124200346) | 1503475 | phosphatidylinositol transfer protein, membrane-associated 2, opposite strand 2 [Source:MGI Symbol;Acc:MGI:3840147] | n.a |
| [Pitpnm2os1](http://www.ensembl.org/mus_musculus/Gene/Summary?db=core;g=ENSMUSG00000090220) | [124229725](http://www.ensembl.org/mus_musculus/contigview?chr=5&vc_start=124229725&vc_end=124237137) | [124237137](http://www.ensembl.org/mus_musculus/contigview?chr=5&vc_start=124229725&vc_end=124237137) | 1538293 | phosphatidylinositol transfer protein, membrane-associated 2, opposite strand 1 [Source:MGI Symbol;Acc:MGI:1923177] | n.a |
| [Mphosph9](http://www.ensembl.org/mus_musculus/Gene/Summary?db=core;g=ENSMUSG00000038126) | [124250959](http://www.ensembl.org/mus_musculus/contigview?chr=5&vc_start=124250959&vc_end=124327972) | [124327972](http://www.ensembl.org/mus_musculus/contigview?chr=5&vc_start=124250959&vc_end=124327972) | 1559527 | M-phase phosphoprotein 9 [Source:MGI Symbol;Acc:MGI:2443138] | [Mphosph9Gt(OST104880)Lex](http://www.informatics.jax.org/allele/MGI:4185170) |
| [Cdk2ap1](http://www.ensembl.org/mus_musculus/Gene/Summary?db=core;g=ENSMUSG00000029394) | [124345417](http://www.ensembl.org/mus_musculus/contigview?chr=5&vc_start=124345417&vc_end=124363082) | [124363082](http://www.ensembl.org/mus_musculus/contigview?chr=5&vc_start=124345417&vc_end=124363082) | 1653985 | CDK2 (cyclin-dependent kinase 2)-associated protein 1 [Source:MGI Symbol;Acc:MGI:1202069] | [Cdk2ap1tm1Dtw; Cdk2ap1Gt(D133C05)Wrst; Cdk2ap1Gt(OST35764)Lex](http://www.informatics.jax.org/allele/MGI:3512185) |
| [Sbno1](http://www.ensembl.org/mus_musculus/Gene/Summary?db=core;g=ENSMUSG00000038095) | [124368702](http://www.ensembl.org/mus_musculus/contigview?chr=5&vc_start=124368702&vc_end=124426001) | [124426001](http://www.ensembl.org/mus_musculus/contigview?chr=5&vc_start=124368702&vc_end=124426001) | 1677270 | strawberry notch 1 [Source:MGI Symbol;Acc:MGI:2384298] | [Sbno1Gt(OST114991)Lex](http://www.informatics.jax.org/allele/MGI:4189743) |
| [Kmt5a](http://www.ensembl.org/mus_musculus/Gene/Summary?db=core;g=ENSMUSG00000049327) | [124439930](http://www.ensembl.org/mus_musculus/contigview?chr=5&vc_start=124439930&vc_end=124462308) | [124462308](http://www.ensembl.org/mus_musculus/contigview?chr=5&vc_start=124439930&vc_end=124462308) | 1748498 | lysine methyltransferase 5A [Source:MGI Symbol;Acc:MGI:1915206] | [Kmt5atm1.1Dare; Kmt5aGt(RRB075)Byg; Kmt5aGt(305D01)Cmhd; Kmt5aGt(D060E05)Wrst; Kmt5aGt(OST1973)Lex](http://www.informatics.jax.org/allele/MGI:3842472) |
| [Rilpl2](http://www.ensembl.org/mus_musculus/Gene/Summary?db=core;g=ENSMUSG00000029401) | [124463265](http://www.ensembl.org/mus_musculus/contigview?chr=5&vc_start=124463265&vc_end=124478366) | [124478366](http://www.ensembl.org/mus_musculus/contigview?chr=5&vc_start=124463265&vc_end=124478366) | 1771833 | Rab interacting lysosomal protein-like 2 [Source:MGI Symbol;Acc:MGI:1933112] | [Rilpl2Gt(OST96650)Lex; Rilpl2Gt(450F7)Cmhd](http://www.informatics.jax.org/allele/MGI:4181783) |
| [Snrnp35](http://www.ensembl.org/mus_musculus/Gene/Summary?db=core;g=ENSMUSG00000029402) | [124483134](http://www.ensembl.org/mus_musculus/contigview?chr=5&vc_start=124483134&vc_end=124491124) | [124491124](http://www.ensembl.org/mus_musculus/contigview?chr=5&vc_start=124483134&vc_end=124491124) | 1791702 | small nuclear ribonucleoprotein 35 (U11/U12) [Source:MGI Symbol;Acc:MGI:1923417] | [Snrnp35Gt(D178F04)Wrst; Snrnp35Gt(OST56118)Lex](http://www.informatics.jax.org/allele/MGI:3893047) |
| [Rilpl1](http://www.ensembl.org/mus_musculus/Gene/Summary?db=core;g=ENSMUSG00000029392) | [124493080](http://www.ensembl.org/mus_musculus/contigview?chr=5&vc_start=124493080&vc_end=124531391) | [124531391](http://www.ensembl.org/mus_musculus/contigview?chr=5&vc_start=124493080&vc_end=124531391) | 1801648 | Rab interacting lysosomal protein-like 1 [Source:MGI Symbol;Acc:MGI:1922945] | [Rilpl1Gt(209A4)Cmhd; Rilpl1Gt(OST684)Lex](http://www.informatics.jax.org/allele/MGI:4969420) |
| [Tmed2](http://www.ensembl.org/mus_musculus/Gene/Summary?db=core;g=ENSMUSG00000029390) | [124540695](http://www.ensembl.org/mus_musculus/contigview?chr=5&vc_start=124540695&vc_end=124550506) | [124550506](http://www.ensembl.org/mus_musculus/contigview?chr=5&vc_start=124540695&vc_end=124550506) | 1849263 | transmembrane p24 trafficking protein 2 [Source:MGI Symbol;Acc:MGI:1929269] | [Tmed2Gt(OST78169)Lex](http://www.informatics.jax.org/allele/MGI:4175586) |
| [Ddx55](http://www.ensembl.org/mus_musculus/Gene/Summary?db=core;g=ENSMUSG00000029389) | [124552864](http://www.ensembl.org/mus_musculus/contigview?chr=5&vc_start=124552864&vc_end=124569660) | [124569660](http://www.ensembl.org/mus_musculus/contigview?chr=5&vc_start=124552864&vc_end=124569660) | 1861432 | DEAD (Asp-Glu-Ala-Asp) box polypeptide 55 [Source:MGI Symbol;Acc:MGI:1915098] | [Ddx55Gt(CSH561)Byg; Ddx55Gt(OST406009)Lex](http://www.informatics.jax.org/allele/MGI:3878538) |
| [Eif2b1](http://www.ensembl.org/mus_musculus/Gene/Summary?db=core;g=ENSMUSG00000029388) | [124570213](http://www.ensembl.org/mus_musculus/contigview?chr=5&vc_start=124570213&vc_end=124579131) | [124579131](http://www.ensembl.org/mus_musculus/contigview?chr=5&vc_start=124570213&vc_end=124579131) | 1878781 | eukaryotic translation initiation factor 2B, subunit 1 (alpha) [Source:MGI Symbol;Acc:MGI:2384802] | [Eif2b1Gt(OST132125)Lex](http://www.informatics.jax.org/allele/MGI:4198203) |
| [Gtf2h3](http://www.ensembl.org/mus_musculus/Gene/Summary?db=core;g=ENSMUSG00000029387) | [124579140](http://www.ensembl.org/mus_musculus/contigview?chr=5&vc_start=124579140&vc_end=124597680) | [124597680](http://www.ensembl.org/mus_musculus/contigview?chr=5&vc_start=124579140&vc_end=124597680) | 1887708 | general transcription factor IIH, polypeptide 3 [Source:MGI Symbol;Acc:MGI:1277143] | [Gtf2h3Gt(D144D12)Wrst; Gtf2h3Gt(RRG412)Byg](http://www.informatics.jax.org/allele/MGI:3892030) |
| [Tctn2](http://www.ensembl.org/mus_musculus/Gene/Summary?db=core;g=ENSMUSG00000029386) | [124598749](http://www.ensembl.org/mus_musculus/contigview?chr=5&vc_start=124598749&vc_end=124627738) | [124627738](http://www.ensembl.org/mus_musculus/contigview?chr=5&vc_start=124598749&vc_end=124627738) | 1907317 | tectonic family member 2 [Source:MGI Symbol;Acc:MGI:1915228] | [Tctn2tm1.1Reit; Tctn2Gt(OST378011)Lex](http://www.informatics.jax.org/allele/MGI:5292130) |
| [Atp6v0a2](http://www.ensembl.org/mus_musculus/Gene/Summary?db=core;g=ENSMUSG00000038023) | [124628576](http://www.ensembl.org/mus_musculus/contigview?chr=5&vc_start=124628576&vc_end=124724455) | [124724455](http://www.ensembl.org/mus_musculus/contigview?chr=5&vc_start=124628576&vc_end=124724455) | 1937144 | ATPase, H+ transporting, lysosomal V0 subunit A2 [Source:MGI Symbol;Acc:MGI:104855] | n.a |
