## Supplementary Figures for "A *P2rx7* passenger mutation affects the vitality and function of immune cells in P2X4ko and other transgenic mice"

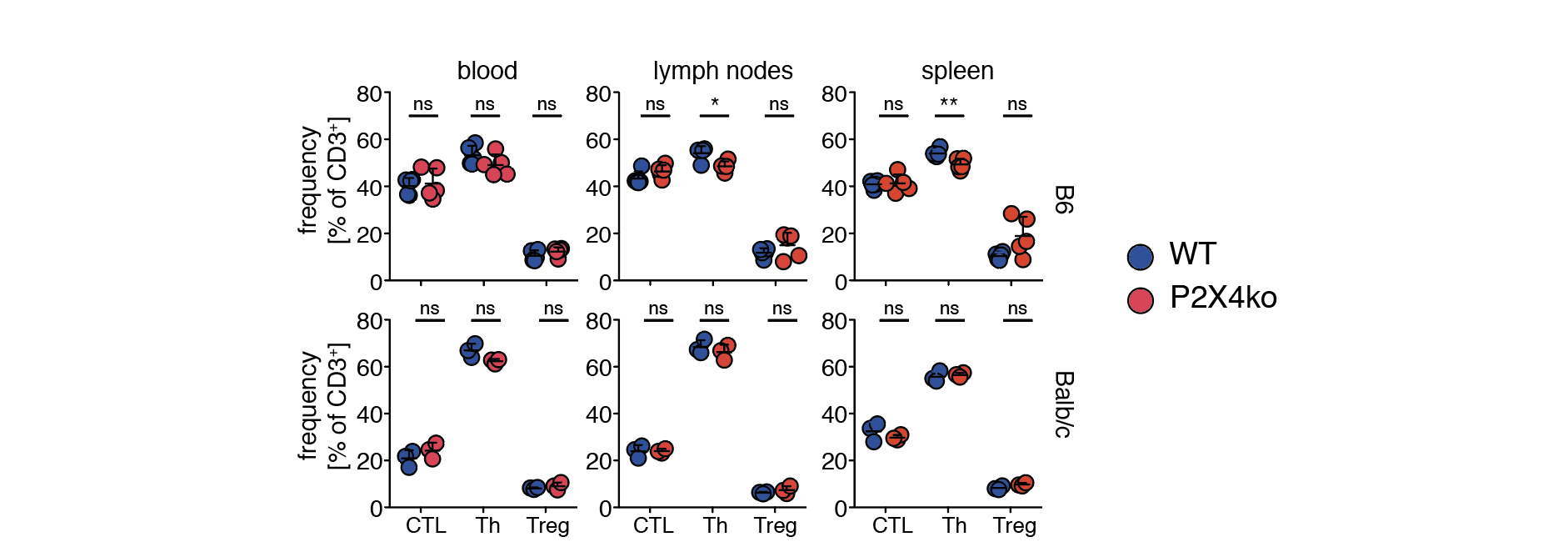

**Supplementary figure S1: T cell frequencies are comparable among P2X4ko and WT mice.** Frequencies of CTL, Th and Treg (n = 3-5) in relation to all CD3^+^ T cells was determined in blood, peripheral lymph nodes and spleen of B6 and Balb/c P2X4ko (red) and WT mice (blue). Statistical comparison of two groups was performed by using the student’s t test (p <  0.05 = * / p < 0.01 = ** / p < 0.001 = ***, ns = no significant).

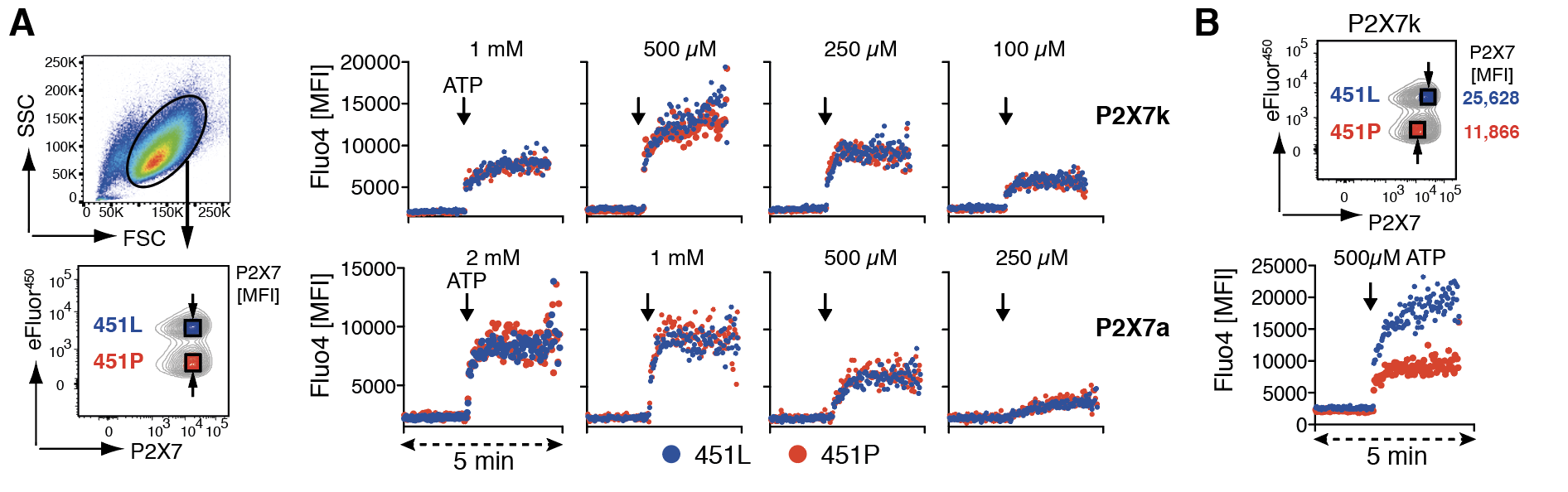

**Supplementary figure S2: The intensity of the calcium influx depends on the expression level of P2X7 . (A)** HEK cells stably transfected with P2X7 451L or 451P (P2X7k or P2X7a splice variant) were distinguished by eFluor^450^ labeling and P2X7 expression level were determined by co-staining with an anti-P2X7 antibody (clone RH23A44). Comparative P2X7 451L/P HEK cell analyses were adjusted for P2X7 expression levels by creating a gating region with a comparable P2X7 mean fluorescence intensity (MFI). For the analyses, mixed HEK cells were loaded with Fluo4 and measured in a real-time flow cytometry assay with 2min baseline recording followed by 3 min ATP stimulation (100µM – 2mM). Calcium influx was measured by increase in Fluo4 MFI. **(B)** Analysis of the 500µM ATP P2X7k sample was repeated with a skewed adjustment of P2X7 expression.

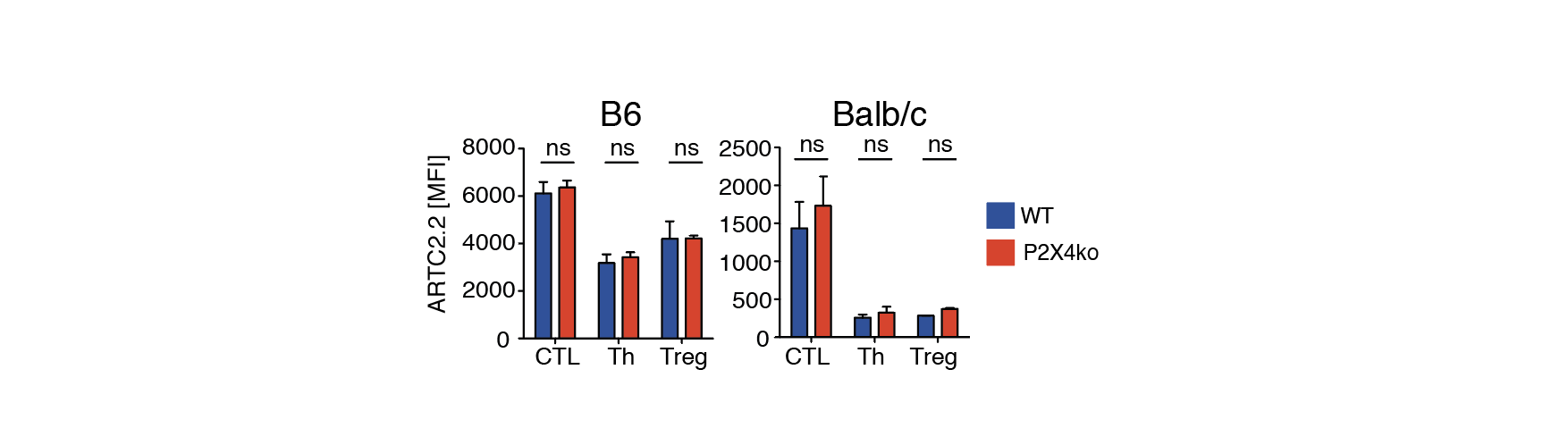

**Supplementary figure S3: ARTC2.2 expression is comparable on P2X4ko and WT T cells.** Flow cytometric analyses of cell surface ARTC2.2 expression on CTL, Th and Treg of WT and P2X4ko mice on the B6 and Balb/c background. The mean fluorescence intensity (MFI) of ARTC2 on the different T cell populations from WT and P2X4ko mice (n = 3) was compared.

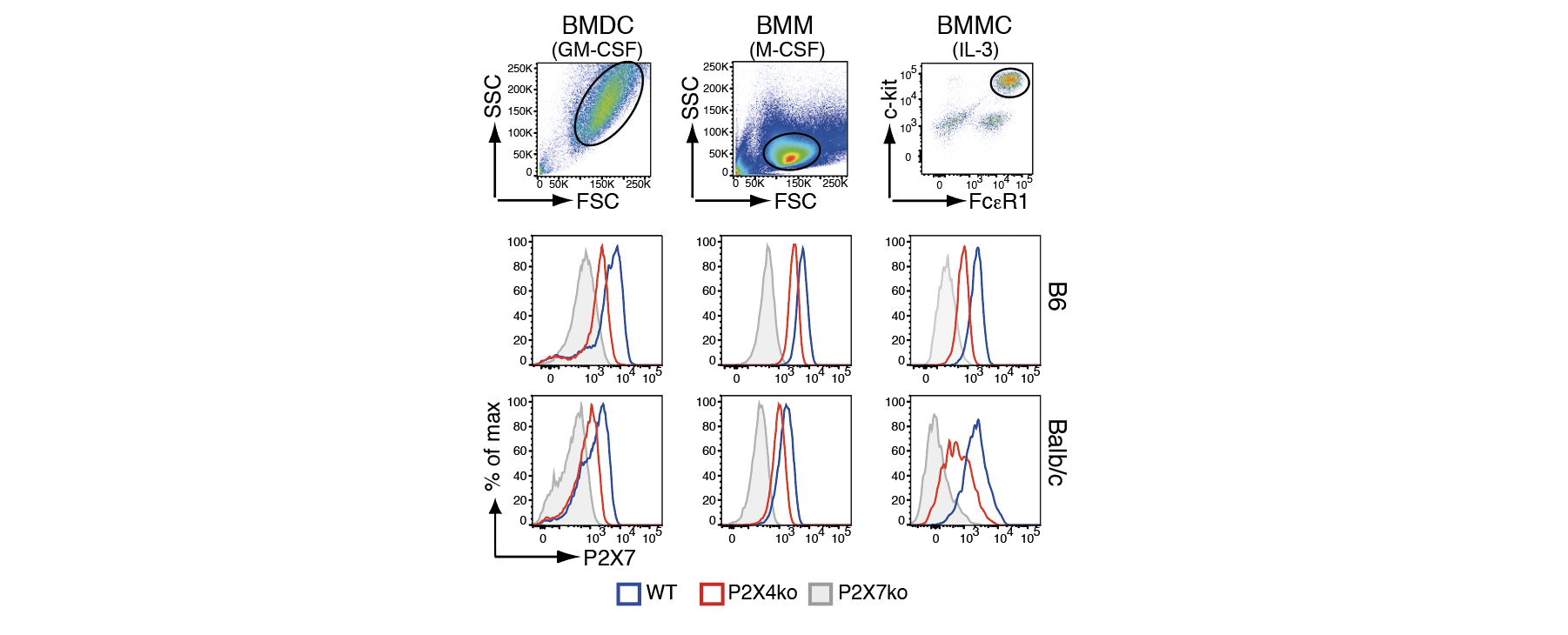

**Supplementary figure 4: Innate immune cells cultured from bone marrow show reduced P2X7 expression.** P2X7 expression was analyzed on bone marrow-derived dendritic cells (BMDC), macrophages (BMM) and mast cells (BMMC) of B6 and Balb/c WT and P2X4ko mice.

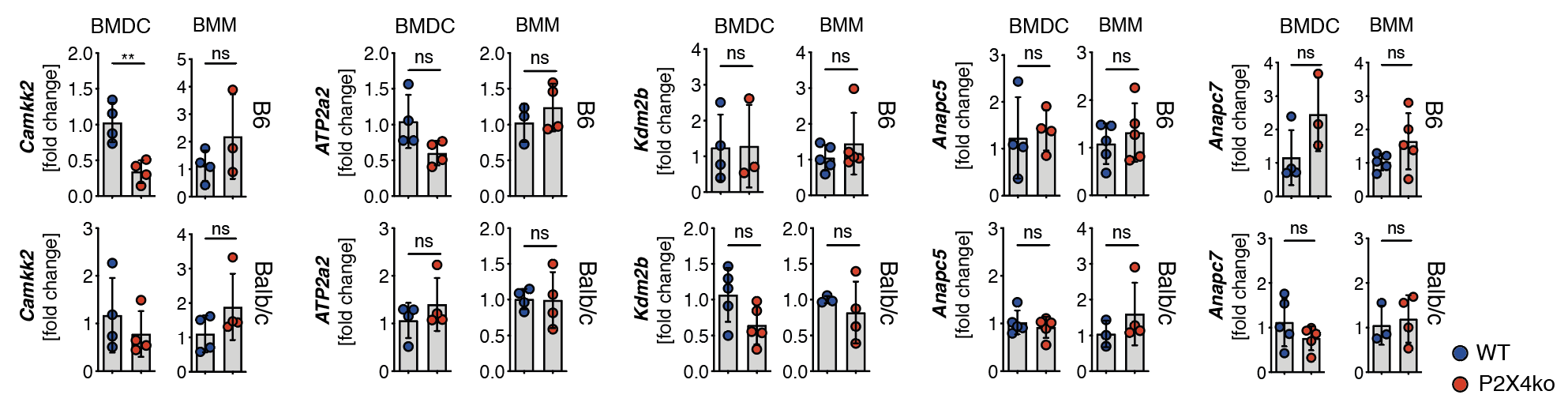

**Supplementary figure 5: mRNA expression of *P2rx4* neighboring genes in innate immune cells of P2X4ko mice.** The expression of the *P2rx4* neighboring genes *Camkk2, ATP2a2, Kdm2b, Anapc5* and *Anapc7* was analyzed in bone marrow-derived dendritic cells (BMDC) and macrophages (BMM) of B6 and Balb/c WT and P2X4ko mice (n = 3-5). Statistical comparison of two groups was performed by using the student’s t test (p <  0.05 = * / p < 0.01 = ** / p < 0.001 = ***, ns = no significant).
